# Evaluation of M2-like macrophage enrichment after diffuse traumatic brain injury through transient interleukin-4 expression from engineered mesenchymal stromal cells

**DOI:** 10.1101/2020.01.28.918441

**Authors:** S. F. Enam, S. R. Kader, N. Bodkin, J. G. Lyon, M. Calhoun, C. Azrak, P. M. Tiwari, D. Vanover, H. Wang, P. J. Santangelo, R. V. Bellamkonda

## Abstract

Appropriately modulating inflammation after traumatic brain injury (TBI) may prevent disabilities for the millions of those inflicted annually. In TBI, cellular mediators of inflammation, including macrophages and microglia, possess a range of phenotypes relevant for an immunomodulatory therapeutic approach. It is thought that early phenotypic modulation of these cells will have a cascading healing effect. In fact, an anti-inflammatory, “M2-like” macrophage phenotype after TBI has been associated with neurogenesis, axonal regeneration, and improved white matter integrity. There already exists clinical trials seeking an M2-like bias through mesenchymal stem/stromal cells (MSCs). However, MSCs do not endogenously synthesize key signals that induce robust M2-like phenotypes such as Interleukin-4 (IL-4). To enrich M2-like macrophages in a clinically relevant manner, we augmented MSCs to transiently express IL-4 via synthetic IL-4 mRNA. We observed that these IL-4 expressing MSCs indeed induce a robust M2-like macrophage phenotype and promote anti-inflammatory gene expression in a modified TBI model of closed head injury. However, here we demonstrate that acute enrichment of M2-like macrophages did not translate to improved functional or histological outcomes. This suggests that an acute enrichment of M2-like macrophages cannot solely orchestrate the neurogenesis, axonal regeneration, and improved WMI after diffuse TBI. To further understand whether dysfunctional pathways underlie the lack of therapeutic effect, we report transcriptomic analysis of injured and treated brains. Through this, we discovered that inflammation persists in spite of acute enrichment of M2-like macrophages in the brain. Last, we comment on our modified TBI model, behavioral studies, and propose that IL-4 expressing MSCs may also have relevance in other cavitary diseases or in improving biomaterial integration into tissues.

## A. INTRODUCTION

Traumatic brain injury (TBI) places a calamitous toll on individuals and the health system. TBI accounts for 138 deaths every day and 2.5 million emergency room visits, hospitalizations, or deaths annually (1). It makes up one-third of all “Unintentional Injuries”, the leading cause of mortality between the ages 1 and 44. This has economic ramifications on the order of $60-80 billion per year (2, 3). TBI is divided into mild, moderate and severe categories. While mild TBI is typically thought to result in a transient loss of function, studies are elucidating persisting complications (4). Severe TBI results in permanent morbidity or mortality in all patients (5).

However, the permanent morbidity and mortality are seldom due to the primary insult and instead result from numerous injury-associated sequelae. These include: inflammation, excitotoxicity, accumulation of reactive oxygen species, apoptosis of injured cells, ischemia, edema, and blood-brain-barrier (BBB) disruption (5). Soon after TBI, damaged cells release distress signals, ‘alarmins’, consisting of intracellular components and various cytokines (6). Some alarmins reach the bloodstream to recruit neutrophils and monocytes that can morph into macrophages in the parenchyma. Other alarmins activate microglia to clear debris and activate astrocytes to reestablish barriers. At first these mechanisms are neuroprotective but over weeks or months they become maladaptive. In mammals, the microglia and macrophages stay persistently skewed towards pro-inflammatory, M1-like phenotypes (7, 8). They self-amplify by releasing more inflammatory cytokines and can remain at injury site for weeks and months (9). The M1-like response in lower-order animals, like zebrafish, is abrogated by a robust ‘anti-inflammatory’ M2-like response within a week, enabling them to regenerate a transected spinal cord (10, 11). Cohorts of M1 and M2 macrophages were first identified *in vitro* and in other injury conditions (12), but the phenotypes can exist concurrently after TBI (13). Additionally, it is now well understood that macrophages and *in vivo* encompass a phenotype across an M1-M2 spectrum. Thus, we use the term M2-like to suggest a phenotype similar to M2 macrophages discovered previously. These M2-like macrophages are, at least indirectly, associated with improved biological and functional recovery in the mammalian peripheral and central nervous system after injury (7,14–18). They are involved in angiogenesis, neurogenesis, axonal regeneration, and improved white matter integrity (19, 20). Therapies that enhance an M2-like phenotype after TBI, often improve molecular markers, edema, white matter, and functional outcomes (21–30) (31–36). However, the peak M2-like response 5 days after injury is still overwhelmed by M1-like macrophages and the desirable response recedes by day 7 (37, 38). Shifting the balance towards endogenous, ‘anti-inflammatory’, M2-like macrophages is thus a viable goal for TBI but requires a clinically relevant strategy. Stem cells are a promising means to promote an anti-inflammatory response after TBI and to possibly modulate macrophage phenotypes (39–44). In fact, a handful of different bone-marrow derived stem cells (BMSCs) are in at least 8 clinical trials (39, 40). These cells include multi-potent adult progenitor cells (MAPCs) and mesenchymal stem/stromal cells (MSCs). Phase I trials have determined that harvesting BMSCs from a patient and delivering them is logistically feasible and safe (40). Encouragingly, at least one Phase I study has reported decreased neural tissue loss and an improvement in a few clinical outcomes at 6 months. Phase II trials are underway in which they compare doses of BMSCs to reduce neuroinflammation after TBI in adults and children. For TBI, one of the more frequently studied stem cells is the MSC (39). Like many other stem cells, they exert their effects not through differentiation but via secretion of cytokines and growth factors, their ‘secretome’. However, while many pre-clinical studies demonstrate their ability to temper inflammation, skepticism towards MSCs persists (40).

Part of the skepticism towards MSC therapy is in the absence of certain growth factors and cytokines in the MSC secretome (40,45–49). One absent cytokine, that also promotes an M2 macrophage phenotype, is Interleukin-4 (IL-4)(46,50,51). IL-4 is a small, 16 kDa protein that is classically secreted by and activates Th2 helper cells. However, when it binds to its cognate receptor IL-4Rα, it initiates STAT6-based transcription and expression of M2 phenotypic proteins on macrophages. *In vivo*, IL-4 promotes M2-like macrophage phenotypes after stroke (52, 53), spinal cord injury (SCI) (54–56), and peripheral nerve injury (PNI) (17, 57) at doses ranging from 250-500 ng. IL-4 stimulates astrocytes to secrete growth factors, and promotes microglia to express M2 phenotypic markers as well (58). Unfortunately, IL-4 levels do not increase in humans or mice after TBI (59, 60). In rats, however, there is a modest IL-4 elevation in the first 24 hours, but this subsides within 3 days (61). Delivery of IL-4 after TBI remains unpublished although it is actively being explored (1I01BX003377-01) (62). Altogether, this suggests that augmenting the MSC secretome with IL-4 could bias macrophages after TBI to a reparative, M2-like, phenotype.

MSCs have previously been genetically altered to express desirable proteins. For rodent models of CNS injury, MSCs have been modified to overexpress bone-derived neurotrophic factor (BDNF) (63), superoxide dismutase 2 (SOD2) (64), interleukin-13 (IL-13) (65–67), or interleukin-10 (IL-10) (47, 49). While most of these modifications have improved molecular parameters in each study, improved functional outcomes are only occasionally observed. BDNF overexpression mildly improved outcomes in one functional SCI assessment (63). SOD2 overexpression demonstrated improved motor coordination at one timepoint tested after focal TBI (64). IL-13 overexpression promotes an M2-like phenotype, but it has only been observed to improve some functional assessments in an SCI model and failed to rescue function in stroke and epilepsy models (65–67). IL-10 overexpression from MSCs did not reduce TBI lesion volume or improve most functional assessments (47, 49). The one functional test that improved (foot faults) and a decreased histological presence of reactive astrocytes was not significantly different from wild-type MSCs. Genetic modification of MSCs to express IL-4 can be achieved via plasmid DNA transfection, viral transduction of DNA, or through synthetic mRNA. MSCs have previously been transduced to express IL-4 and consequently promoted a Th2 response in a model of autoimmune encephalitis (51). Modification of MSCs in this manner requires DNA that encodes IL-4 to enter the nucleus and then induces prolonged or permanent expression of IL-4. Persistent IL-4 expression, although initially therapeutic, is undesirable as it promotes a chronic M2-like macrophage presence which results in fibrosis (23, 68). This persistent overexpression can be advantageous outside the nervous system; implanted MSCs over-expressing IL-4 increase bone mineral density (50). However, in the CNS, fibrosis is deleterious. Viral transduction also comes with the risk of undesired genomic integration and activation of oncogenes. In contrast, synthetic mRNA only needs to reach ribosomes in the cytosol, thus enabling rapid expression and a short persistence of 1-4 days (47). This approach could counter an M1-like response faster and allow the injured site to return to homeostasis. Lower-order animals that regenerate their CNS show a similar dynamic: pro-inflammatory followed by anti-inflammatory peaks that all subside to homeostasis within a couple weeks (10, 11).

Delivering recombinant IL-4 alongside MSCs is another strategy though it can also have drawbacks. As mentioned, IL-4 alone has been delivered after stroke, SCI, and PNI. This method may have rescued functional deficits in IL-4 knockout mice suffering stroke (52) and reports in SCI are conflicting (56, 69). This ambiguity could be because recombinant protein delivery has yet to fully address the challenges of short half-lives (70, 71), suboptimal efficacy (72), and immunogenicity (73, 74). Cytokines possess short half-lives due to rapid circulation and clearance *in vivo* and this is exacerbated by increased interstitial flow after TBI. In mouse models, IL-4 half-life can be less than 20 minutes (51, 75). To counter this, multi-dosing or sustained release strategies need to be employed. A lack of post-translational modifications (PTMs) and the presence of microstructural abnormalities can further worsen half-life, reduce efficacy, and increase immunogenicity. These challenges can be addressed by inducing harvested MSCs to express IL-4 via nucleic acids as unsullied IL-4 can be synthesized with endogenous PTMs.

In the present study, we transfect MSCs with synthetic IL-4 mRNA to transiently induce IL-4 expression. We hypothesize that these modified MSCs will promote M2-like macrophage mediated healing in a modified closed head injury (CHI) mouse model of TBI. Through our model, we investigate *in vivo* macrophage polarization, examine behavioral outcomes, scrutinize cytokine and gene expression, probe histological alterations in tissue and white matter tracts, and explore transcriptomic dysregulation.

## B. RESULTS

### a. Transfection with Viromer Red and synthetic IL-4 mRNA generates IL-4 expression

As MSCs do not endogenously produce IL-4, introduction of IL-4 genetic information is required. To reliably introduce synthetic IL-4 mRNA into MSCs, transfection agents exist that enable the mRNA to reach the cytosol without degradation. Here, we first obtained MSCs (Cyagen; harvested from C57BL/6N mice; later *in vivo* studies are conducted in this mouse strain) that expressed putative MSC markers (76), although there was an absence of CD105. The cells retained this marker expression at passages 9 and 12 as well (S. Fig. 1) but most experiments were conducted between passages 7 and 9. Next, synthetic mRNA encoding Green Fluorescent Protein (GFP) was combined with one of three transfection agents (jetPEI, Lipofectamine Messenger Max, or Viromer Red) and incubated for 24 hours with MSCs following manufacturer protocol for each agent at 0.5x and 1x their recommended dose. The amount of synthetic mRNA (250 or 500 ng) was equivalent between the transfection agents. Of the three, Viromer Red induced the greatest mean fluorescence intensity (MFI) in a flow cytometer and was thus chosen for future transfections (Fig. 1A). Next, we tested Viromer Red toxicity by complexing it with synthetic IL-4 mRNA and incubating four doses with MSCs for 24 hours. Regardless of the dose, less than 5% of all MSCs were seen to be dead or dying via a commercial live/dead cell assay (Fig 1B). Simultaneously, IL-4 ELISA confirmed that at least 150 ng of IL-4 was expressed into the media (1 mL) (Fig 1C). Without transfection of synthetic IL-4 mRNA, MSCs did not endogenously produce IL-4 (Fig 1D). In another study, we characterized transfection-induced expression over a 48-hour period and observed that IL-4 synthesis begins within the first 2-4 hours and the majority is synthesized within the first 24 hours (Fig 1E). These studies confirm that MSCs do not endogenously produce IL-4, and transfection with Viromer Red and synthetic IL-4 mRNA can generate IL-4 expression for at least 24 hours with acceptably low toxicity.

**Figure 1:**
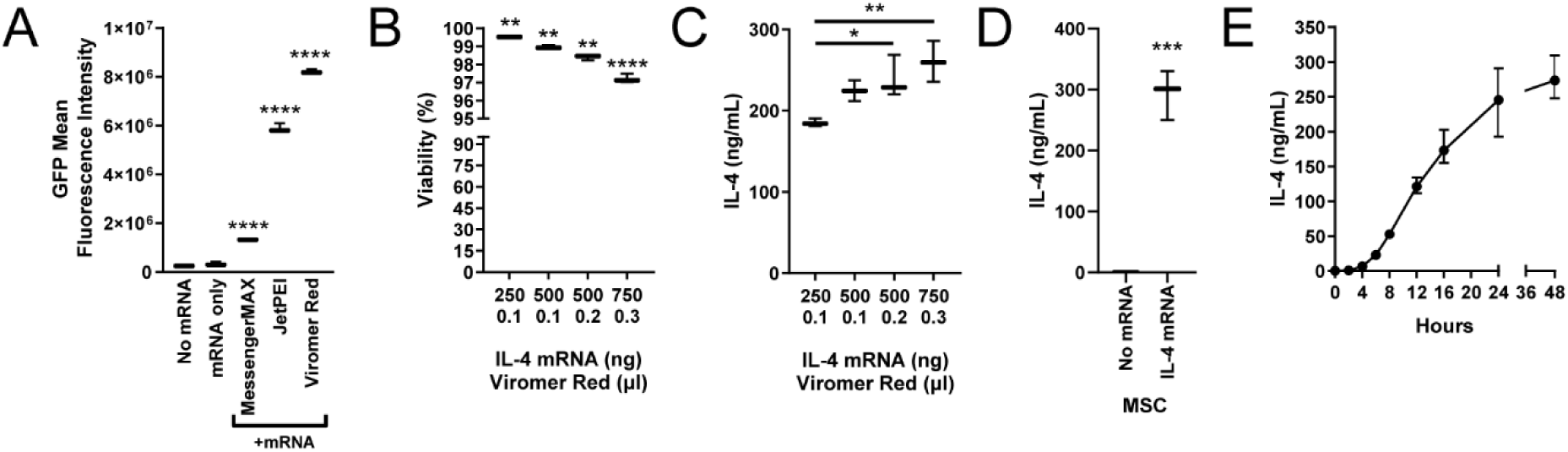
Transfection of MSCs and protein expression. **(A)** GFP fluorescence intensity after MSCs are transfected with naked GFP mRNA or complexed with one of three vectors. **(B)** Viability of MSCs with increasing dosage of synthetic IL-4 mRNA complexed with Viromer Red. **(C)** Concentration of IL-4 synthesized in media with IL-4 mRNA transfected MSCs. **(D)** MSCs do not endogenously synthesize IL-4 and require transfection with IL-4 mRNA. **(E)** Concentration of IL-4 synthesized over 48 hours from IL-4 mRNA transfected MSCs. Mean ± SD, one-way ANOVAs with Tukey’s post-hoc test and Student’s t-test; \**p* < 0.05, \*\**p* < 0.01, \*\*\**p* < 0.001, \*\*\*\**p* < 0.0001.

**Supplementary Figure 1:**
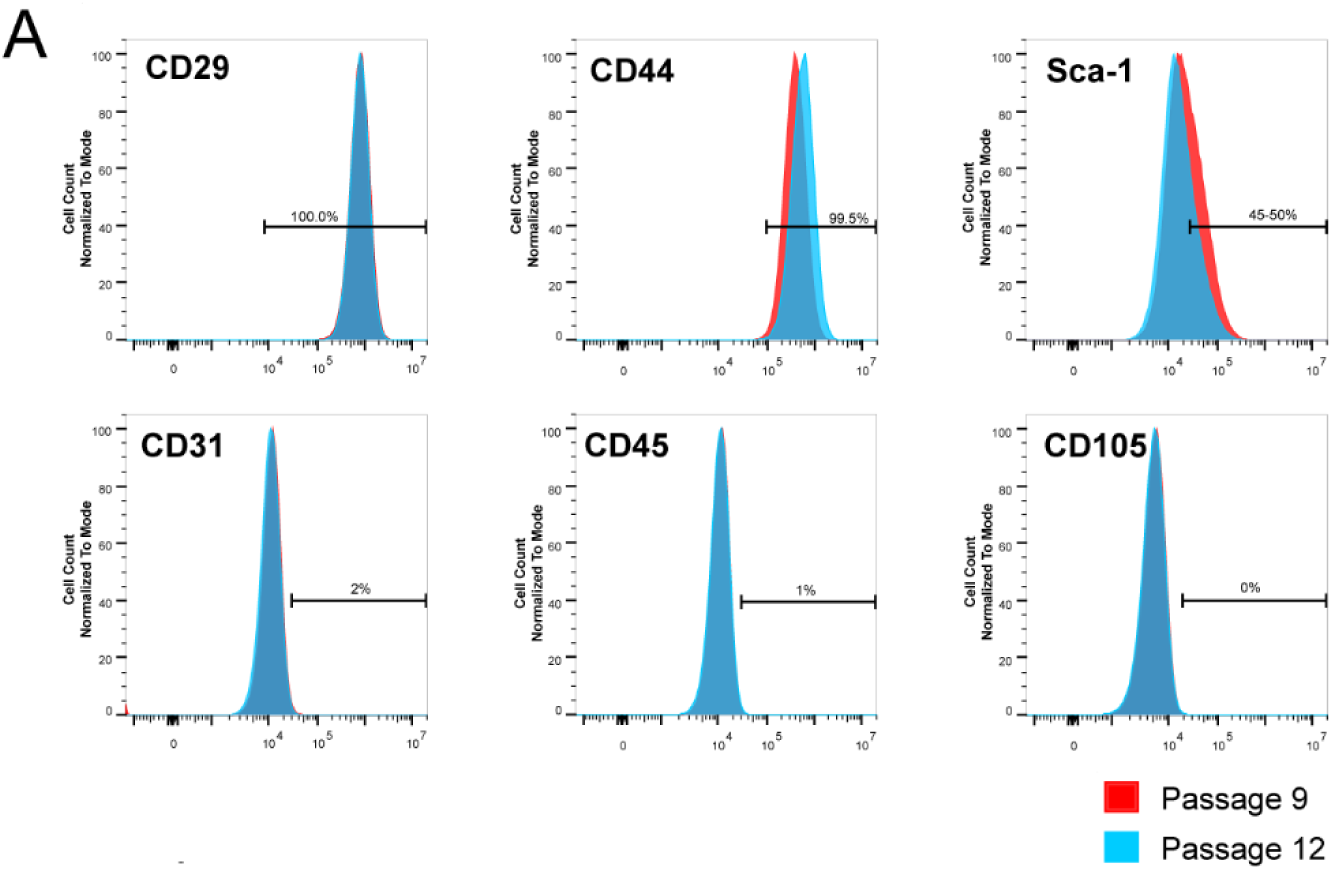
MSC cell-surface markers. **(A)** MSCs (Cyagen) used in this study expressed CD29, CD44, and Sca-1 while remaining negative for CD31 and CD45 in at least two late passage numbers.

### b. Expressed IL-4 induces an M2-like phenotype *in vitro*

Next, we set out to determine whether IL-4 secreted from transfected MSCs was functional. IL-4 is a potent stimulator of the M2 cell-surface marker, CD206, on macrophages. We set up four groups: plain media, media conditioned by naïve MSCs (MSC CM), MSC CM supplemented with 100 ng of recombinant IL-4 (rIL-4 CM), and media conditioned with IL-4 mRNA transfected MSCs (synthesizing ∼80-100 ng expressed IL-4; eIL-4 CM) (Fig. 2A). We incubated the media with cultures of J774A.1 macrophages for 24 hours after which time they were harvested, stained, and analyzed in a flow cytometer. As hypothesized, macrophages treated with either rIL-4 CM or eIL-4 CM demonstrated significantly greater expression of CD206 (Fig. 2B). As expected, CD163, an alternative M2-like marker (typically stimulated by IL-10), was not upregulated by IL-4 (Fig. 2C). Control experiments demonstrated that macrophage cell morphology and division rate was unaffected by media formulated for MSCs (S. Fig. 2A-C). To rule out the possibility of expressed IL-4 having an autocrine effect on MSCs, the MSCs were also treated with 100 ng of IL-4 for 24 hours, thoroughly washed, and then CM was collected another 24 hours later. When this CM was added to macrophages, there was no increase in CD206 expression (S. Fig. 2D). These studies demonstrated that IL-4 generated from synthetic IL-4 mRNA-transfected MSCs could induce an M2-like phenotype in macrophages.

**Figure 2:**
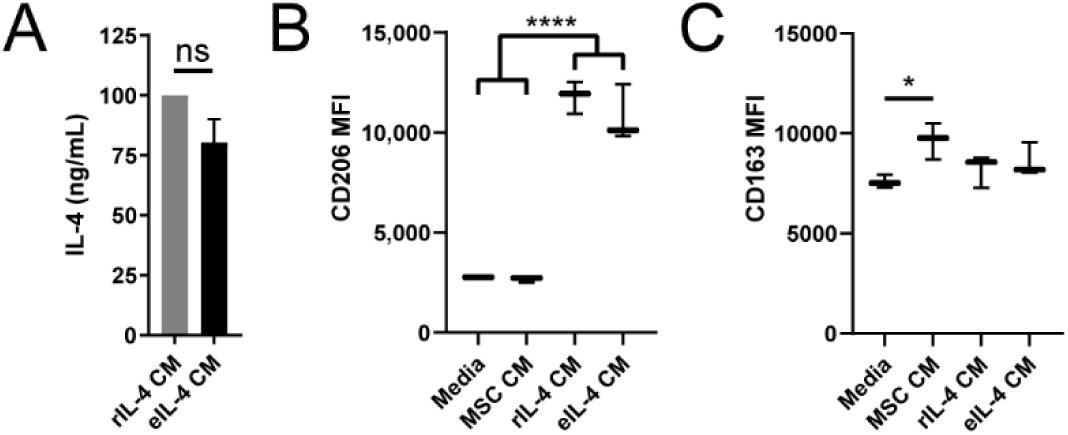
Functionality of IL-4 expressed by IL-4 mRNA transfected MSCs. **(A)** IL-4 concentration in MSC conditioned media (CM) that was either supplemented with 100 ng/mL recombinant IL-4 (rIL-4 CM) or transfected with IL-4 mRNA (eIL-4 CM); ns = not significant. **(B)** Mean Fluorescence Intensity (MFI) of CD206 expression on macrophages after treatment with differently conditioned media. Analyzed in a flow cytometer with a fluorescent CD206 antibody. Both rIL-4 CM and eIL-4 CM were significantly different from Media and naïve MSC CM. **(C)** Mean fluorescence intensity of CD163 expression on macrophages after treatment with differently conditioned media. Mean ± SD, Student’s t-test and one-way ANOVA with Tukey’s post-hoc test; \**p* < 0.05, \*\*\*\**p* < 0.0001.

**Supplementary Figure 2:**
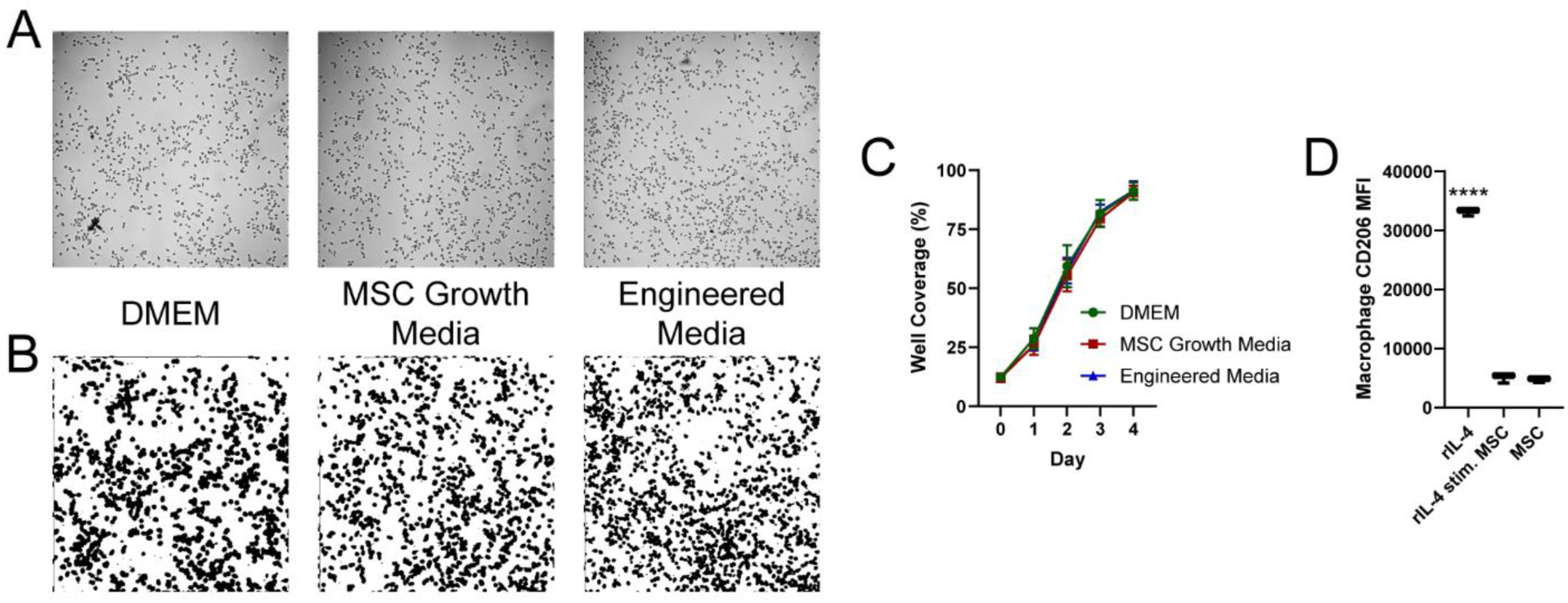
Macrophage and MSC control experiments. **(A)** Brightfield images of macrophages grown in recommended media (Dulbecco’s Modified Eagle Media, DMEM), MSC growth media, or MSC media engineered to match molecular concentrations of DMEM. Each image is 1/16^th^ of total image analyzed per well. **(B)** Masks from automated custom ImageJ quantification script to calculate area of plate covered by cells. **(C)** Growth curves of macrophages in each of the media estimated by coverage of the well, calculated from masks. **(D)** Mean fluorescence intensity of CD206 expression on macrophages after treatment with either 100 ng/mL recombinant IL-4 (rIL-4), conditioned media from MSCs pre-stimulated with 100 ng/mL rIL-4, or conditioned media from naïve MSCs. Mean ± SD, one- or two-way ANOVA; \*\*\*\**p* < 0.0001.

### c. Transient MSC transfection is a viable strategy for *in vivo* experiments

Due to the transience of IL-4 expression after mRNA transfection, it was critical to characterize expression considering the logistics of *in vivo* delivery. First, we set out to determine the ideal incubation time of IL-4 mRNA/Viromer Red complexes with MSCs to synthesize maximal IL-4. MSCs were incubated with the complexes for varying durations (0-24 hours). After this period, the cells were washed twice with PBS and left to express IL-4 in fresh media for 24 hours. Quantification of IL-4 via ELISA demonstrated that 8-12 hours of transfection elicited the greatest IL-4 production over the subsequent 24 hours (Fig. 3A). Next, as multiple *in vivo* cell injections were expected to take 4-6 hours, MSC survival on ice was assessed through trypan blue exclusion. Here, MSCs demonstrated >95% viability over at least six hours on ice (Fig. 3B). We also hypothesized that viability in media would be superior to that in PBS, however we did not observe a significant difference. All *in vivo* injections were thus conducted in PBS to reduce variability. Curiously, media sampled from these transfected MSCs while on ice demonstrated minimal IL-4 expression (Fig. 3C). Then, regardless of how long the MSCs were on ice (0-6 hours), the amount of IL-4 produced over the next 24 hours in a 37°C incubator was equivalent (Fig. 3C). Finally, to scale for *in vivo* delivery, 300,000 cells per well were transfected for 10 hours with either 1 or 2 µg of IL-4 mRNA and a proportional amount of Viromer Red. Media was collected after 10 hours prior to harvesting cells. After cells were harvested, 150,000 cells per well were re-plated. Media was collected again after 24 hours. The 300,000 cells transfected with 2 µg induced at least 120 ng of IL-4 prior to harvest. The harvested 150,000 cells then produced another ∼135 ng of IL-4 over 24 hours (Fig 3D). This transfection protocol was chosen for all *in vivo* experiments; MSCs transfected thusly are henceforth denoted IL-4 MSCs.

**Figure 3:**
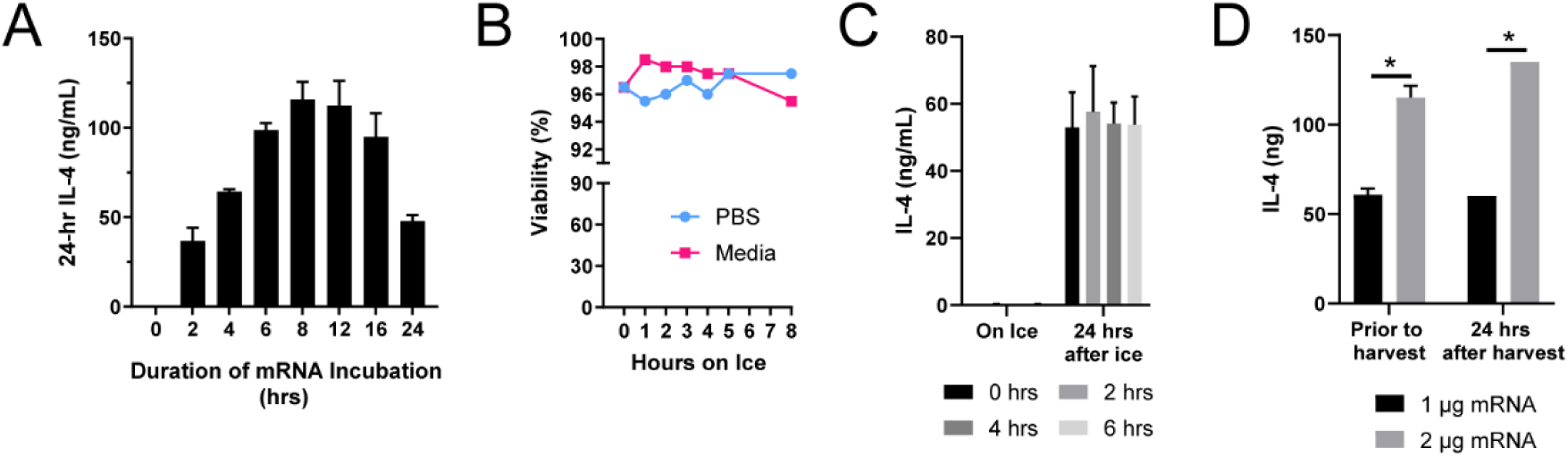
MSC transfection characterization for *in vivo* delivery. **(A)** Concentration of IL-4 in media of MSCs transfected with synthetic IL-4 mRNA complexed with Viromer Red. The complexes were incubated with the MSCs for varying amounts of time (0-24 hours). Then, the media was sampled 24 hours after each timepoint for IL-4 quantification via ELISA. **(B)** Viability of MSCs while on ice. No significant differences observed. **(C)** Concentration of IL-4 synthesized while transfected and harvested MSCs were kept on ice (0-6 hrs) and then over 24 hours after re-plating and kept at 37°C. **(D)** Amount of IL-4 expressed by 300,000 MSCs transfected for 10 hours (‘prior to harvest’) and then a sample of 150,000 MSCs 24 hours after. Mean ± SD, two-way ANOVA comparing between groups with Tukey’s post-hoc; \**p* < 0.05.

### d. IL-4 MSCs induce an M2-like phenotype *in vivo* after brain injury

To assess whether IL-4 MSCs can promote an M2-like phenotype of macrophages *in vivo*, IL-4 MSCs were injected in a TBI model of closed-head-injury (CHI) in mice. This model is similar to a model developed by *Webster, et al.,* but induces a motor deficit (described in methods and in Fig. 5) (77). Five days after injury, we injected either PBS, MSCs, MSCs expressing Luciferase (Luc MSCs), recombinant IL-4 (rIL-4), or IL-4 MSCs into the left hippocampus. For groups containing MSCs, 150,000 cells were injected, a number based on prior studies (42). Brains were harvested from euthanized mice two days after injection and leukocytes isolated from the ipsilateral hemisphere were analyzed with flow cytometry (Fig. 4A). First, we observed an increase in total macrophage (CD45^high^ F4/80^+^) number for most groups (Fig. 4B). Of these macrophages, IL-4 MSCs induced 60-80% to an M2-like phenotype (CD206^+^, CD86^-^) (Fig. 4C). This was significantly higher than PBS (<10%), MSCs alone (20%), or Luc MSCs (<10%). In comparison, an injection of 250 ng of rIL-4 induced CD206 expression in 10-25% of macrophages (Fig 4C). IL-4 is known to induce proliferation of M2-like macrophages (78) and we observed that there were more CD206^+^ macrophages in the IL-4 MSC group compared to all other groups (Fig. 4D). We also calculated an M2:M1 ratio: [CD206^+^] / [CD206^-^, CD86^+^], a metric shown to be predictive of neural regeneration in a peripheral nerve injury model (17, 57). IL-4 MSCs dramatically increased the M2:M1 ratio compared to all other groups (Fig. 4E). While MSCs and rIL4 had a higher percentage of M2-like macrophages (Fig. 4C), the M2:M1 ratio showed no difference compared to control (Fig. 4E). An increase in the mRNA/Viromer Red transfection dose to MSCs greatly increased the number of M2-like macrophages (S. Fig. 3 A and C) while pushing the polarization only mildly higher to 80% (S. Fig. 3B). Interestingly, the M2:M1 ratio was lower with this higher dose suggesting a low dose is more desirable (S. Fig. 3D). Next, as CHI is diffuse, we required a delivery strategy that would affect the macrophages in both the ipsilateral (left) and contralateral (right) hemispheres. We hypothesized that delivering IL-4 MSCs into a lateral ventricle, instead of the hippocampus, would induce greater CD206^+^ macrophage expression in the contralateral hemisphere. Contrary to this hypothesis, the number of macrophages, mean percentage M2-like polarization, and number of M2-like macrophages in both ipsi- and contra-lateral hemispheres was lower with an intraventricular versus an intrahippocampal injection (S. Fig. 3E-G). Thus, all subsequent injections were made into the left hippocampus. One week after injection, the number of macrophages had fallen in all groups (Fig. 4F). The M2-like phenotype (CD206^+^, CD86^-^) dropped to ∼37% of macrophages one week after IL-4 MSC injection (Fig. 4G). Intriguingly, ∼30% of macrophages in the PBS group retained an M2-like phenotype while ∼20% of macrophages in the MSC only group did. However, the number of CD206^+^ macrophages was still significantly greater in the IL-4 MSC group compared to the others (Fig. 4H). A modified M2:M1 ratio was calculated as some groups did not possess any CD206^-^, CD86^+^ cells. This ratio was again significantly higher in the IL-4 MSC group compared to PBS and MSCs alone (Fig. 4I). These data demonstrate that IL-4 MSCs can increase the number of M2-like macrophages for at least up to 1 week after delivery and comprise 60-80% of all macrophages 2 days after injection. Although one week after intervention similar percentages of the M2-like phenotype exist regardless of treatment, a desired M2:M1 ratio persists with IL-4 MSCs.

**Figure 4:**
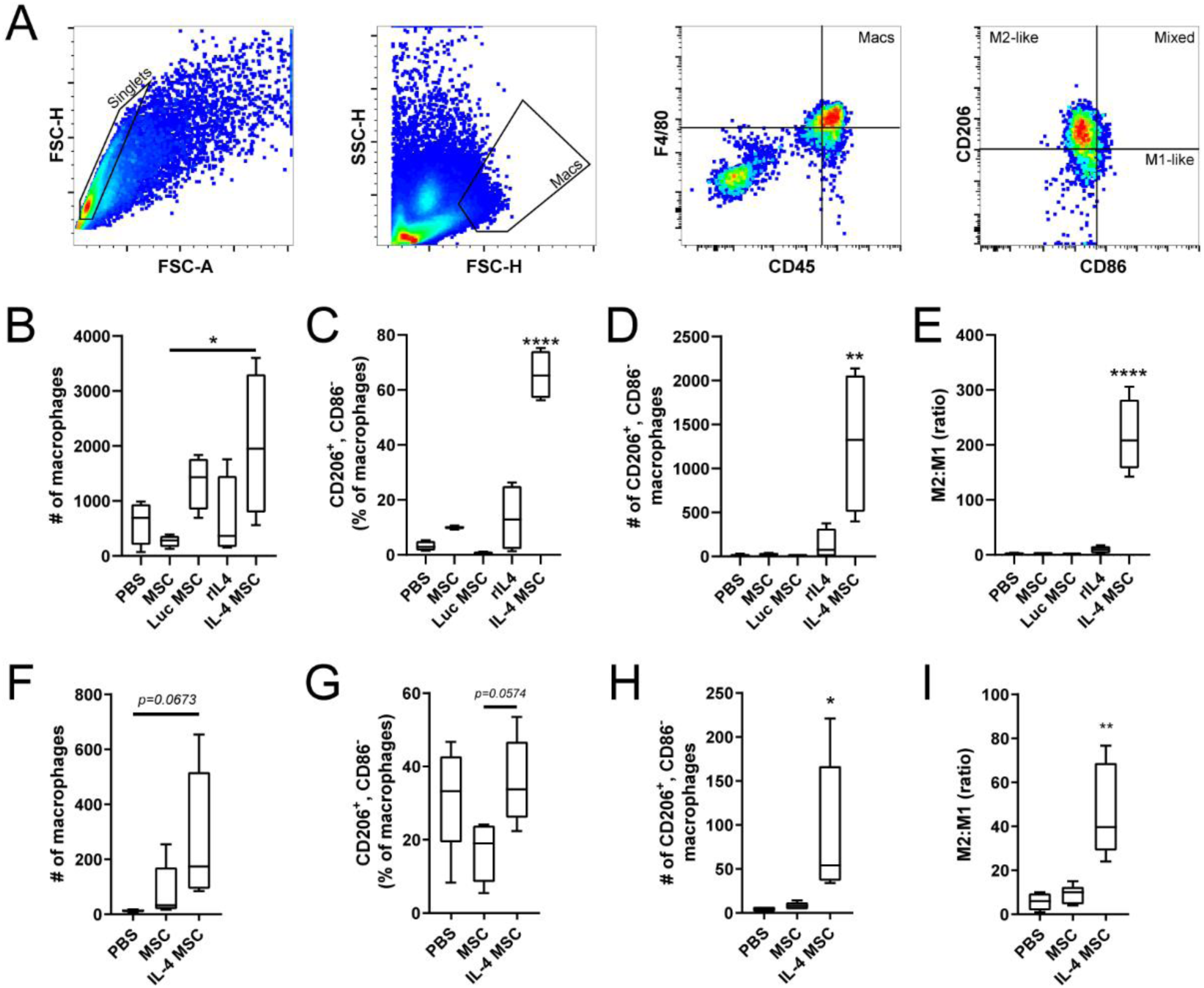
Macrophage polarization after closed head injury and treatments. **(A)** Flow cytometry gating to identify macrophages (CD45^hi^, F4/80^+^) and M2-like macrophages (CD206^+^, CD86^-^). Labeled gates produce the subsequent image. **(B) – (E) Macrophage analysis 1 week after injury, 2 days after treatment; *n* = 4, N = 20. (B)** Number of total macrophages in the ipsilateral hemisphere for each treatment group. **(C)** Percentage of total macrophages that possess the M2-like phenotype (CD206^+^, CD86^-^). **(D)** Number of M2-like macrophages in each treatment group. **(E)** Ratio of M2-like to M1-like macrophages calculated by [CD206^+^, CD86^+/-^] / [CD206^-^, CD86^+^]. **(F) – (I) Macrophage analysis 12 days after injury, 1 week after treatment; *n* = 5, N = 15. (F)** Number of total macrophages. **(G)** Percentage of total macrophages that possess the M2-like phenotype (CD206^+^, CD86^-^). **(H)** Number of M2-like macrophages. **(I)** Modified M2:M1 ratio. In samples where there were ‘0’ M1-like cells, this was changed to ‘1’ to enable calculation of ratio. Mean ± SD, one-way ANOVA used for each graph with Tukey’s post-hoc; \**p* < 0.05, \*\**p* < 0.01, \*\*\*\**p* < 0.0001.

**Supplementary Figure 3:**
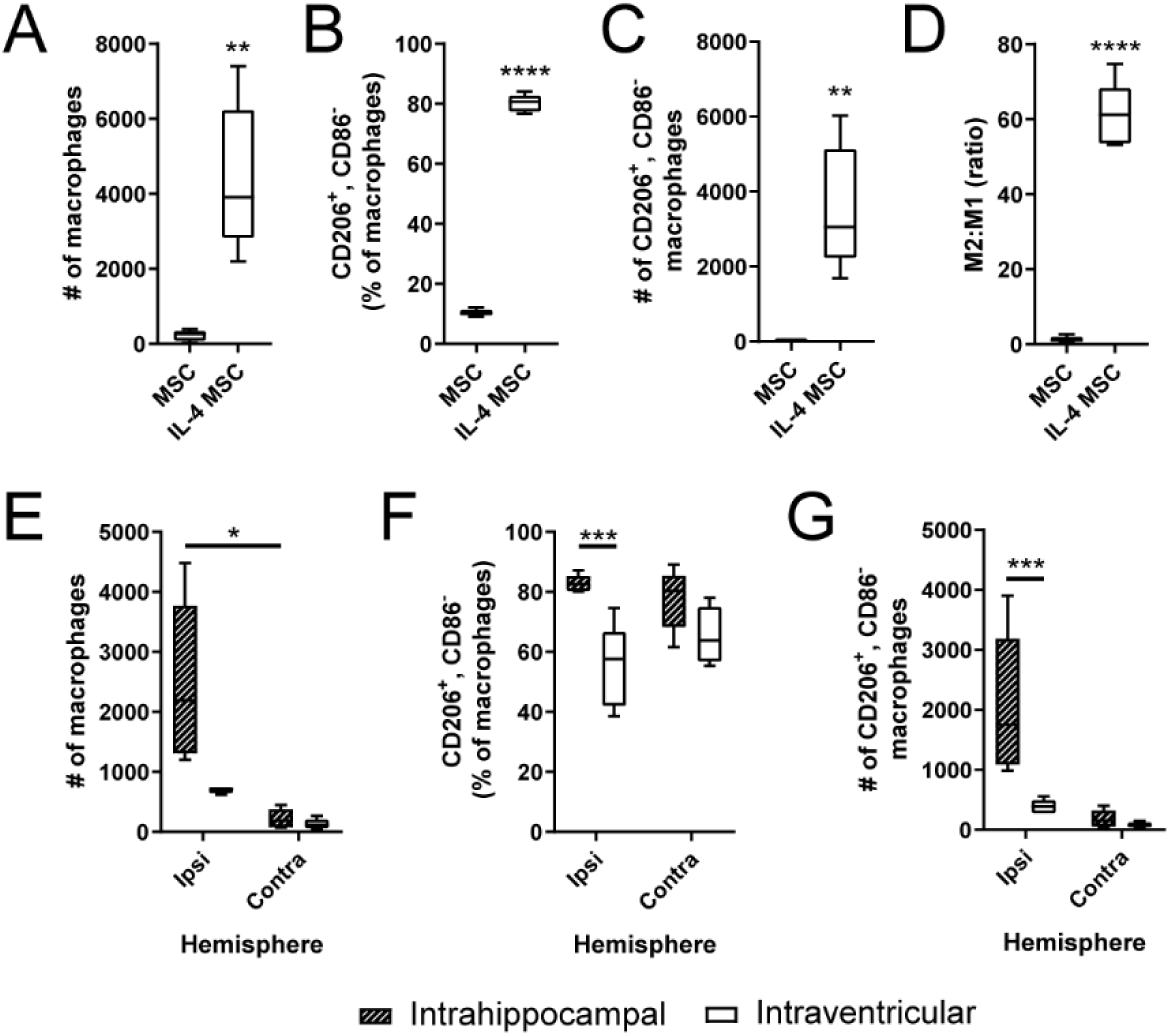
Macrophage polarization after closed head injury and treatment alterations through flow cytometry. (A) – (D) Macrophage analysis 1 week after injury, 2 days after treatment with IL-4 MSCs producing greater IL-4; *n* = 5, N = 10. **(A)** Number of total macrophages in the ipsilateral hemisphere for each treatment group. **(B)** Percentage of all macrophages that possess M2-like phenotype. **(C)** Number of macrophages with M2-like phenotype. **(D)** Ratio of M2 to M2-like macrophages. **(E) – (G) Macrophage analysis 1 week after injury, 2 days after treatment delivered into the left hippocampus or left lateral ventricle; *n* = 5, N = 10. (E)** Total number of macrophages in the ipsilateral and contralateral hemispheres after either delivery modality. **(F)** Percent of total macrophages that possess an M2-like phenotype. **(G)** Number of M2-like macrophages in either hemisphere. Mean ± SD, Student’s t-tests or two-way ANOVAs with post-hoc Tukey’s; \**p* < 0.05, \*\**p* < 0.01, \*\*\**p* < 0.001, \*\*\*\**p* < 0.0001.

### e. IL-4 MSCs do not rescue acute injury-induced motor dysfunction

To assess whether the dramatic polarization and robust increase of CD206^+^ macrophages improved functional outcomes after CHI, injured mice with and without IL-4 MSC treatment underwent two behavioral tests. Motor coordination/performance was evaluated via persistence on an accelerating Rotarod for at least 1 week. Spatial memory was assessed via time spent solving the Morris Water Maze (MWM), 3-4 weeks after injury. Our modified CHI model (with no injections) (Fig. 5A) was first compared to mice receiving a sham procedure: all experimental steps aside for the actual injury and injection. Injured mice demonstrated poorer Rotarod performance in the first two weeks (Fig. 5B) and poor spatial memory one month out, compared to the sham group (Fig. 5C). Severity of CHI was assessed via time for the mouse to right itself from a supine position, as previous reports indicate a Righting Time (RT) >10 min to indicate a severe injury (79). However, logistic regression of RT against Day 1 Rotarod performance did not produce a strong correlation (S. Fig. 4A). Neurologic Severity Score (NSS), a more laborious severity assessment, was also unsuccessful in predicting day 1 Rotarod performance (S. Fig. 4B) (80). Next, injured mice received either PBS or IL-4 MSCs five days after injury into the left hippocampus. Rotarod assessment demonstrated no significant differences in performance between the two groups (Fig 5D). A repeat of this experiment demonstrated similar results. MWM assessment demonstrated significant differences between the injured groups and sham, but no difference between IL-4 MSCs and PBS (Fig 5E). At baseline, injured mice who received an intrahippocampal injection took longer to find the hidden platform than sham and those who only received an injury (Fig. 5 C and E). Finally, to test the hypothesis that an earlier change in macrophage phenotype could improve outcomes, we performed intrahippocampal injections 2 days after injury. However, this also did not rescue acute changes in motor coordination (Fig. 5F). MWM was not repeated as Fig. 5E suggests the intrahippocampal injection itself affects performance. These behavioral studies suggested that regardless of when IL-4 MSCs were delivered, poor Rotarod performance could not be rescued.

**Figure 5:**
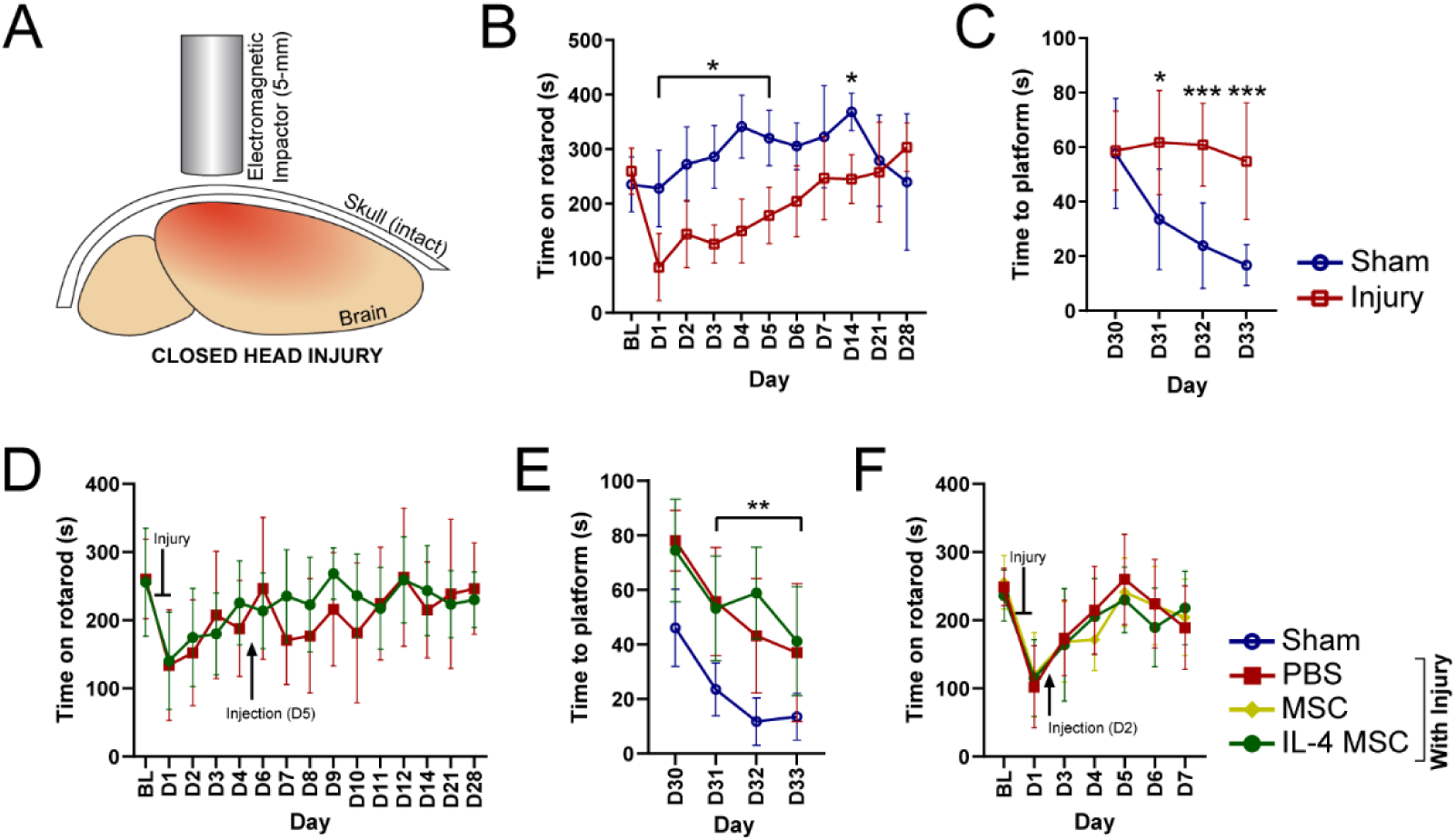
Behavioral analysis after closed head injury and treatment. **(A)** Model representing the direct impact to an intact skull in closed head injury, leading to a diffuse cortical injury. (B), (C) Functional analyses in sham mice (open blue circle) and mice injured on Day 0 (open red square); *n* = 10, N = 20. (B) Time spent running on an accelerating rotarod (4 to 40 RPM). Time was recorded when the mouse fell off or if it grabbed on to the rod for ≥ 2 consecutive spins. **(C)** Time taken to find the hidden glass platform in a Morris Water Maze (MWM) task. (D), (E) Functional analyses in Day 0 injured mice with either day 5 treatment of PBS (closed red square) or IL-4 MSC (closed green circle) (*n* = 10), or sham mice (open blue circle) (*n* = 5); N = 25. (D) Time spent running on an accelerating rotarod. Blunt end indicates point of injury, arrow indicates day of injection. **(E)** Time taken to find the hidden glass platform in a MWM task. (F) Functional analysis in Day 0 injured mice with either day 2 treatment of PBS, IL-4 MSC, or wild-type MSC (closed yellow diamond); *n* = 10, N = 30. (F) Time spent on an accelerating rotarod with markers for day of injury and injection. BL = baseline, D = Day; Graphs show mean ± SD; Repeated Measures ANOVAs with post-hoc Tukey’s in each study; \**p* < 0.05, \*\**p* < 0.01, \*\*\**p* < 0.001.

**Supplementary Figure 4:**
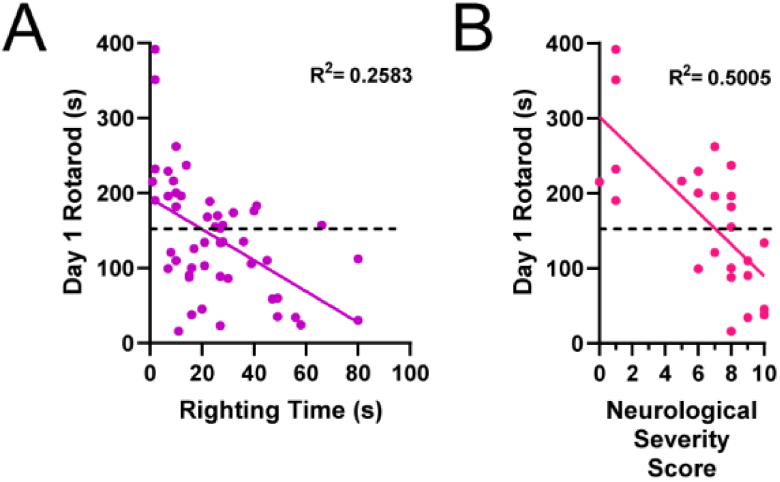
Correlation measures for injury severity. **(A)** Correlation of time for mouse to right itself after injury with day 1 performance on an accelerating Rotarod task. **(B)** Correlation of 10-point Neurological Severity Score with day 1 performance on an accelerating Rotarod task. Analyzed via linear regression; dashed line indicates rotarod performance of 150 s; R^2^ values are reported; correlation was considered to be present if R^2^ > 0.65 and *p* < 0.05.

### f. IL-4 MSCs increase IL-4 in both cerebral hemispheres and anti-inflammatory genes

In light of the conflicting M2-polarization and behavioral data, we next explored how IL-4 MSCs affected inflammatory and anti-inflammatory cytokines and a select panel of genes. Injured mice received PBS, MSCs only, or IL-4 MSCs, and were compared to sham mice (no injury or injection). Multi-plex ELISA demonstrated that cerebral IL-4 levels were higher in the IL-4 MSC group two days after the intervention compared to all other groups (Fig. 6A). This held true in both ipsi- and contralateral hemispheres across the groups. Within the IL-4 MSC group, IL-4 levels were also significantly higher in the ipsilateral hemisphere (0.45 ± 0.34 pg/mg) compared to the contralateral (0.03 ± 0.01 pg/mg) (Fig. 6A). IL-6, a cytokine with a protective role in TBI that is secreted by M2-like macrophages (81), was also significantly higher in the ipsilateral hemispheres of IL-4 MSCs mice compared to sham and PBS (Fig. 6B). There was no difference in IL-6 among the contralateral hemispheres (Fig. 6B). Mean IL-2 was also reduced in the IL-4 MSC group in both ipsi- and contralateral hemispheres but this did not reach statistical significance (Fig. 6C). As anticipated, there were no significant differences in the other five cytokines studied among the four groups (S. Fig. 5) as cytokines tend to normalize to sham levels within a week after injury. A set of 26 genes were investigated 1 and 3 weeks after injury (2 and 16 days after treatment) through Taqman probes and are presented as a heatmap (Fig. 6D). PCR data was analyzed on Applied Biosystems software with global normalization. ΔΔCT was calculated for each gene against all sham mice. One week after injury, genes of two inflammatory cytokines (*il1β, ccl2*) and one anti-inflammatory cytokine (*il1rn*) were significantly upregulated at least >2-fold in all groups (PBS, MSC, IL-4 MSC) compared to sham mice (Fig. 6D and S. Table 1). However, *il1rn* (encoding the IL-1 receptor antagonist) was upregulated 20-25X in PBS and MSC groups while it was upregulated 85X in the IL-4 MSC group (S. Table 1). M2-like genes *mrc1* (encoding CD206) and *cd163* were elevated only in the IL-4 MSC group. Numerous genes assayed were also downregulated in all groups at this 2-day timepoint including growth factors and genes related to axonal regeneration. Three weeks after injury, *il1rn* upregulation had persisted in MSC and IL-4 MSC groups. Genes encoding growth factors and markers of neurogenesis tended to increase in expression at the 3-week timepoint, however most of these did not reach statistical significance. The IL-4 MSC group also had statistically elevated *sox2* (associated with neurogenesis or stem cells) and *casp7* (associated with apoptosis). These data suggest that while IL-4 MSCs acutely increase intracranial IL-4 levels and alters genes associated with inflammation, there is little to no long-term effect on genes associated with neurogenesis or growth factors.

**Figure 6:**
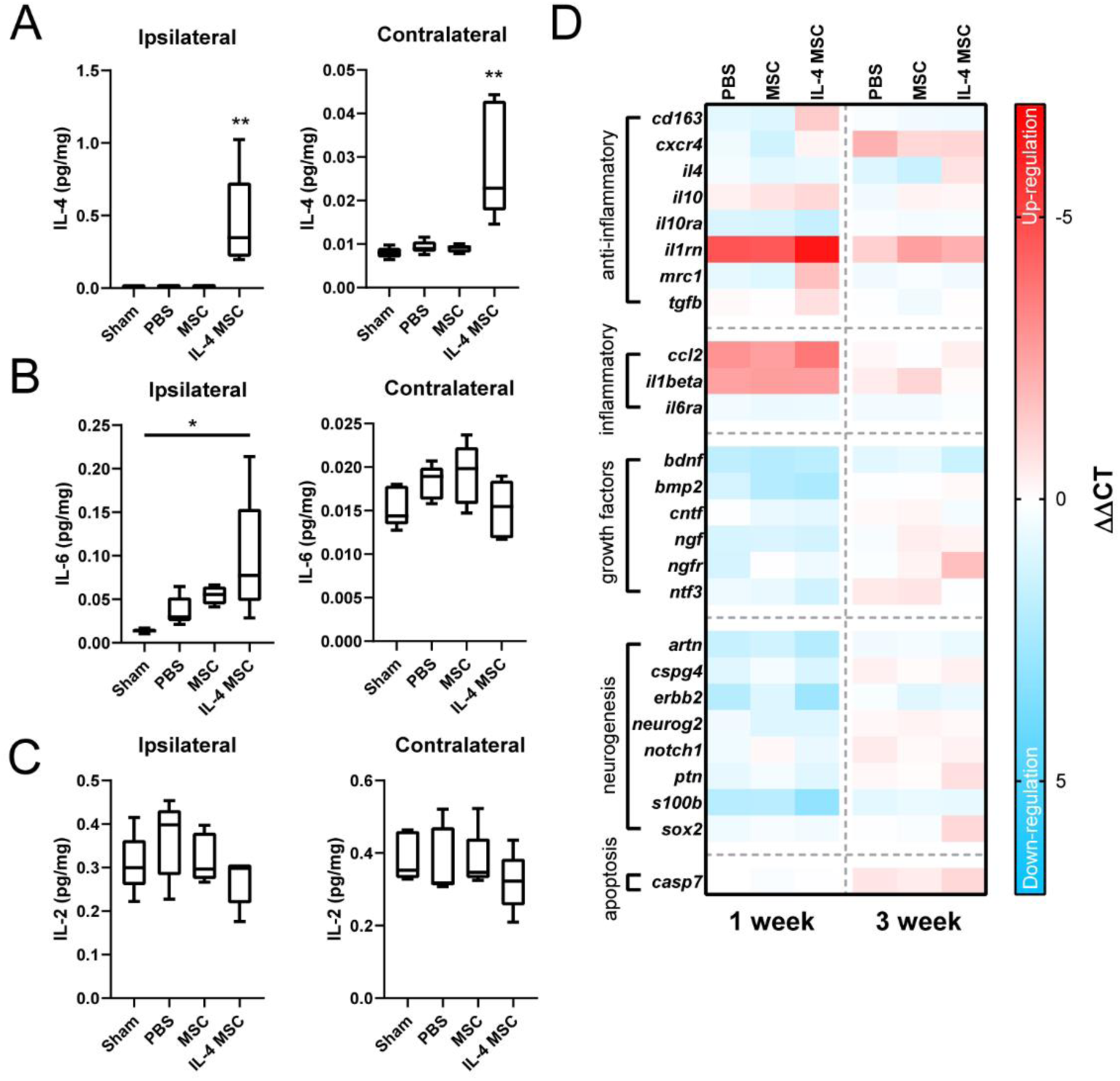
Molecular analysis of *ex vivo* injured and treated brain tissue. (A) – (C) Protein analysis at day 7 of brain from sham mice and mice injured (day 0) and treated on day 5 with either PBS, MSCs, or IL-4 MSCs; *n* = 5, N = 20. (A) Amount of Interleukin-4 (IL-4) in ipsilateral and contralateral cortical samples, normalized to total protein present in the cortical sample, for each mouse group. **(B)** Amount of Interleukin-6 (IL-6) in ipsilateral and contralateral cortical samples, normalized to total protein present in the cortical sample, for each mouse group. **(C)** Amount of Interleukin-2 (IL-2) in ipsilateral and contralateral cortical samples, normalized to total protein present in the cortical sample, for each mouse group. (D) Gene analysis at 1 week or 3 weeks after injury and day 5 treatment with either PBS, MSCs, or IL-4 MSCs (*n* = 5) and sham mice as biological controls (*n* = 10); N = 40. (D) Heatmap of 26 genes demonstrating up- (red) or down- (blue) regulation of genes based on ΔΔCT values. Graphs present mean ± SD; one-way ANOVA carried out for each with a post-hoc Tukey’s comparison; \**p* < 0.05, \*\**p* < 0.01.

**Supplementary Figure 5:**
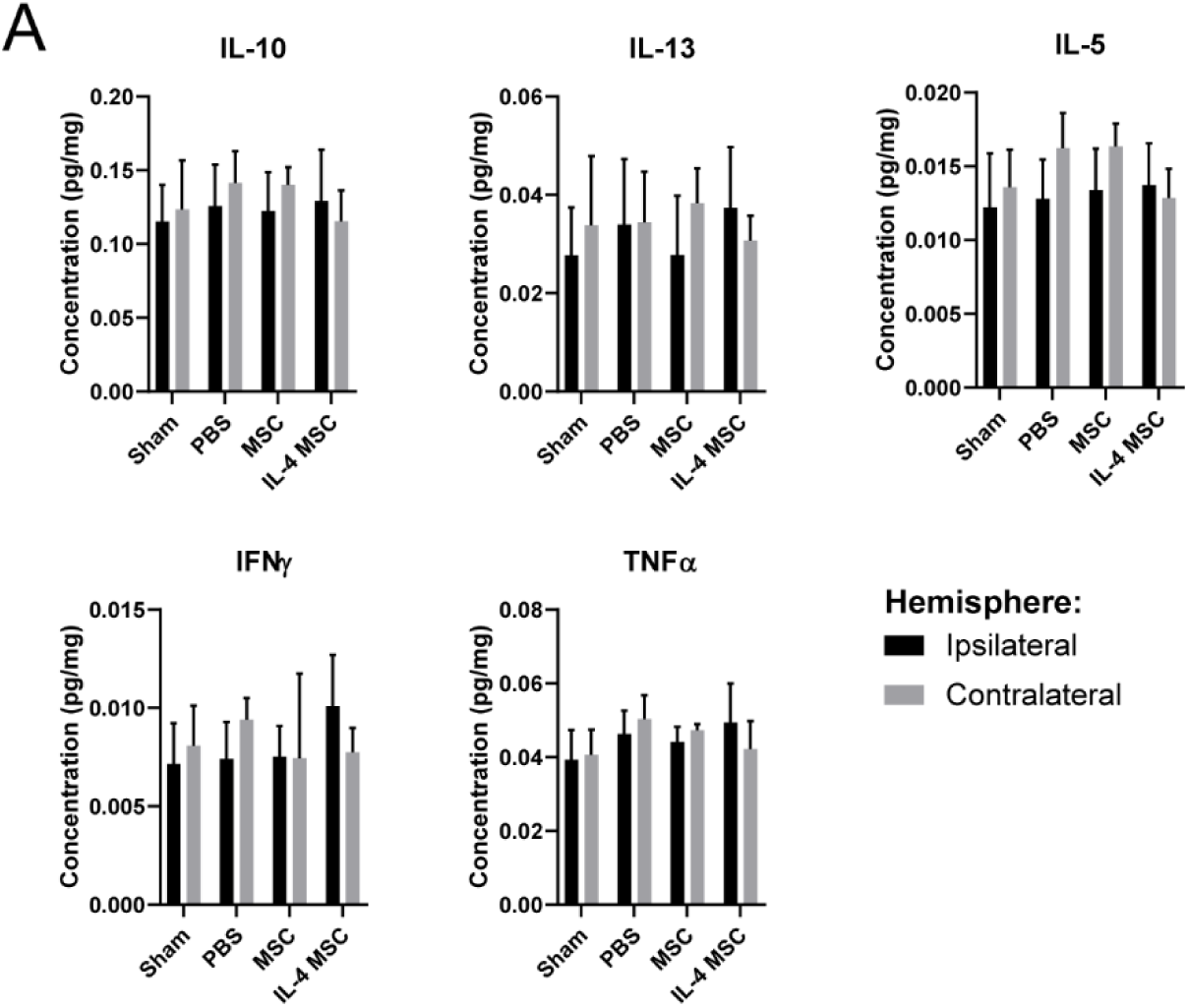
Cytokine analysis of *ex vivo* injured and treated brain tissue at 1 week after injury. Sham mice or injured mice with day 5 treatment of either PBS, MSCs, or IL-4 MSCs; *n* = 5, N = 20. **(A)** Amounts of each cytokine (Interleukin 10, IL-10; Interleukin-13, IL-13; Interleukin-5, IL-5; Interferon-γ, IFNγ; and Tumor Necrosis Factor α, TNFα) per hemisphere normalized to total protein in that hemisphere. Black bars represent the ipsilateral hemisphere while gray bars represent the contralateral hemisphere. Graphs display mean ± SD; two-way ANOVA carried out for each cytokine.

**Supplementary Table 1:**
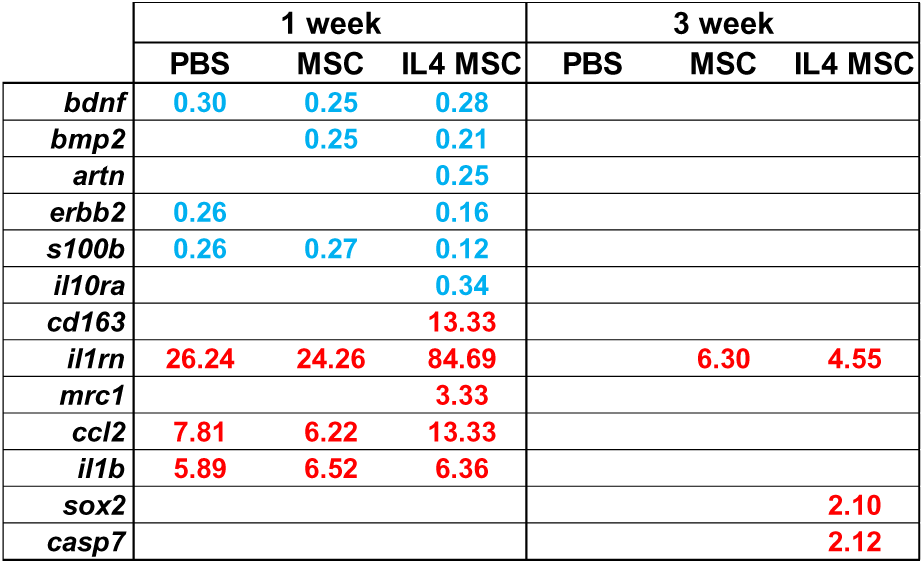
Significantly altered gene expression. The table lists the genes that had a significant (adjusted *p* < 0.05) up- (red) or down- (blue) regulation (log2[fold change] > 2) compared to sham mice at 1 and 3 weeks for each treatment after injury.

### g. IL-4 MSCs increase macrophages but do not reduce astrogliosis or neuronal degeneration

We next scrutinized histological changes due to the absence of behavioral differences in the presence of increased M2-like macrophages. Markers for macrophages, astrogliosis, microglia, and neuronal degeneration were evaluated through immunohistochemistry (IHC) at 1 and 3 weeks after injury (2 and 16 days after intervention). Mice were injured and treated with either PBS, MSCs, or IL-4 MSCs and compared to sham mice. Brain regions analyzed included: ipsilateral and contralateral hippocampi, cortices, and the corpus callosum. Mean fluorescence was quantified through a semi-automated custom ImageJ script (S. Materials). GFAP (a marker for reactive astrocytes) was elevated in all injured groups compared to sham (Fig. 7A) at both time points in both cortices (Fig. 7 B and C). Expression tended to be greater in the injected hemisphere (ipsilateral, left) at 1 week and was also elevated in the ipsilateral hippocampus. However, there was no difference among the treatment groups. CD68 (a marker for macrophages and microglia) and Iba1 (microglia) was absent in the sham group (Fig. 7D) but significantly elevated in all injury groups in both hemispheres (Fig. 7E-G). CD68 expression was greater in the IL-4 MSC group compared to PBS in both the ipsilateral cortex and corpus callosum (Fig. 7 F and G). CD68^+^ cells were also seen across the corpus callosum and in the contralateral hemisphere (Fig. 7G). We also observed dramatic structural alterations in the ipsilateral hippocampus caused by the injection needle (Fig 7 E and G). At the 3-week timepoint, a cluster of cells, likely IL-4 MSCs, was visibly surrounded by macrophages and microglia (Fig. 7H, left). MSCs were absent in the PBS group, but staining nonetheless indicated presence of macrophages and microglia (Fig. 7H, right). Fluoro-Jade C staining demonstrated no neuronal degeneration in sham mice (Fig. 7I) but was detectable in all injured groups at both timepoints (Fig. 7 J and K). Greater staining was visible in the cortices and hippocampus at the 1-week time point (Fig. 7J) than at later timepoints. These histological assessments demonstrate that our CHI model induces a diffuse astrocytosis which is not reduced by IL-4 MSCs, and that CHI also recruits macrophages which is further enhanced by IL-4 MSCs.

**Figure 7:**
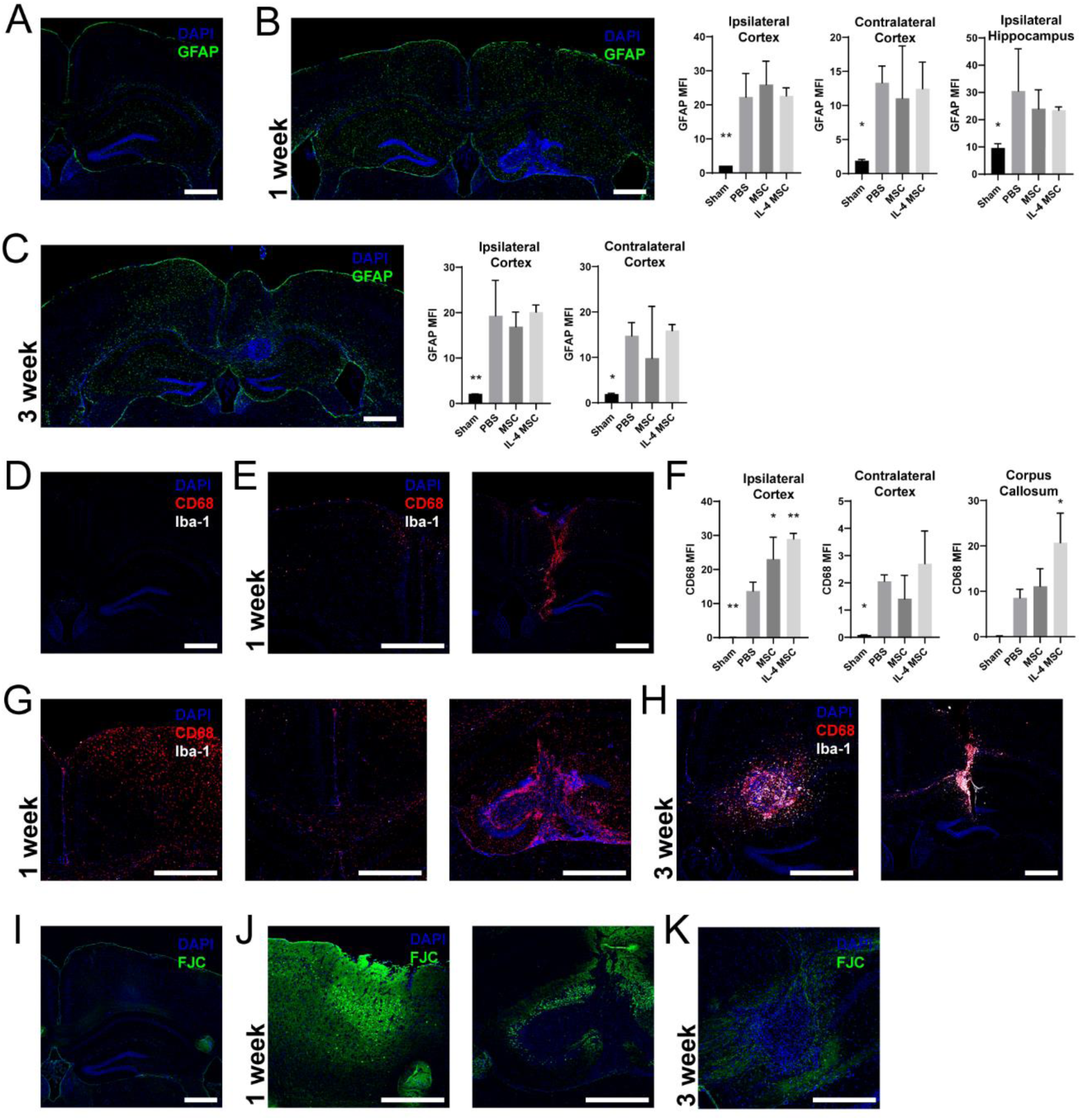
Immunohistochemical analysis of injured and treated brains at 1 and 3 weeks after injury. **(A)** Representative image of mouse brain section from sham group stained for GFAP (green) and DAPI (blue). **(B)** Representative image of mouse brain section from an injured group 1 week after injury. Image demonstrates diffuse reactive astrocytosis in both hemispheres. Graphs show mean fluorescence intensity (MFI) of GFAP in the ipsilateral and contralateral cortices, and the ipsilateral hippocampus for each group. **(C)** Representative image of a mouse brain section from an injured group 3 weeks after injury and stained for GFAP and DAPI. Graphs show GFAP MFI in the cortices for each group at this timepoint. **(D)** Representative image of brain section from the sham mouse group stained for DAPI (blue), CD68 (red), and Iba1 (white). **(E)** Representative sections from an injured mouse brain 1 week after injury who received PBS treatment on Day 5. Left image shows macrophage infiltration in the contralateral cortex. Right image shows the extent of needle damaging the cortex and macrophage infiltration. **(E)** Graphs depict CD68 MFI across the treatment groups in both cortices and the corpus callosum. **(G)** Representative images from an injured mouse brain 1 week after injury who received IL-4 MSCs on Day 5. Left image shows extensive macrophage infiltration in the left cortex. Middle image shows macrophage migration in the corpus callosum. Right image shows needle-induced damage and macrophage infiltration of the hippocampus. **(H)** Representative images from brains 3 weeks after injury. Left image is a section from a mouse treated with IL-4 MSCs. It shows a cluster of cells surrounded by macrophages and microglia. Right image is a section from a mouse treated with PBS. **(I)** Representative image of brain section from sham mouse group stained with DAPI and Fluoro-Jade C. **(J)** Representative images 1 week after brain injury. Left images demonstrates neuronal degeneration in the cortex. Right image shows neuronal degeneration in the hippocampus. **(K)** Representative image 3 weeks after brain injury with less neuronal degeneration than after the first week. Graphs shows mean ± SD; one-way ANOVAs with Dunnett’s multiple comparison against the PBS group. \**p <* 0.05, \*\**p* < 0.01.

### h. IL-4 MSCs do not improve white matter integrity after injury

In brain repair, M2-like macrophages are associated with improved white matter integrity (WMI) (19, 20). To assess whether the M2-like phenotype induced by IL-4 MSCs improved WMI after TBI, a subset of injured mice treated with PBS or IL-4 MSCs and a subset of sham mice were scanned 35 days after injury with diffusion-weighted magnetic resonance imaging (Fig. 8 A and B). The images were mapped to a dataset to automatically orient and identify white matter regions of interest (Fig. 8 A and B). Values of fractional anisotropy (FA) and diffusivity (axial, radial, and mean) were obtained for each region. White matter tracts are anisotropic and when subjected to trauma, this anisotropy is reduced (82, 83). Thus, we hypothesized that our injury would also disrupt white matter tracts and thus reduce FA while increasing diffusivity. Compared to sham, FA values were lower in the PBS and IL-4 MSC groups in all white matter tracts however no difference was observed between the two treatments (Fig. 8C). FA was particularly reduced in the brachium of the superior colliculus, corpus callosum, and optic tracts (Fig. 8C) of both injured and treated groups. While there were trends in diffusivity measures, differences between groups were not statistically significant. This MRI study demonstrated that while injury reduced WMI, IL-4 MSCs did not improve WMI one month after delivery.

**Figure 8:**
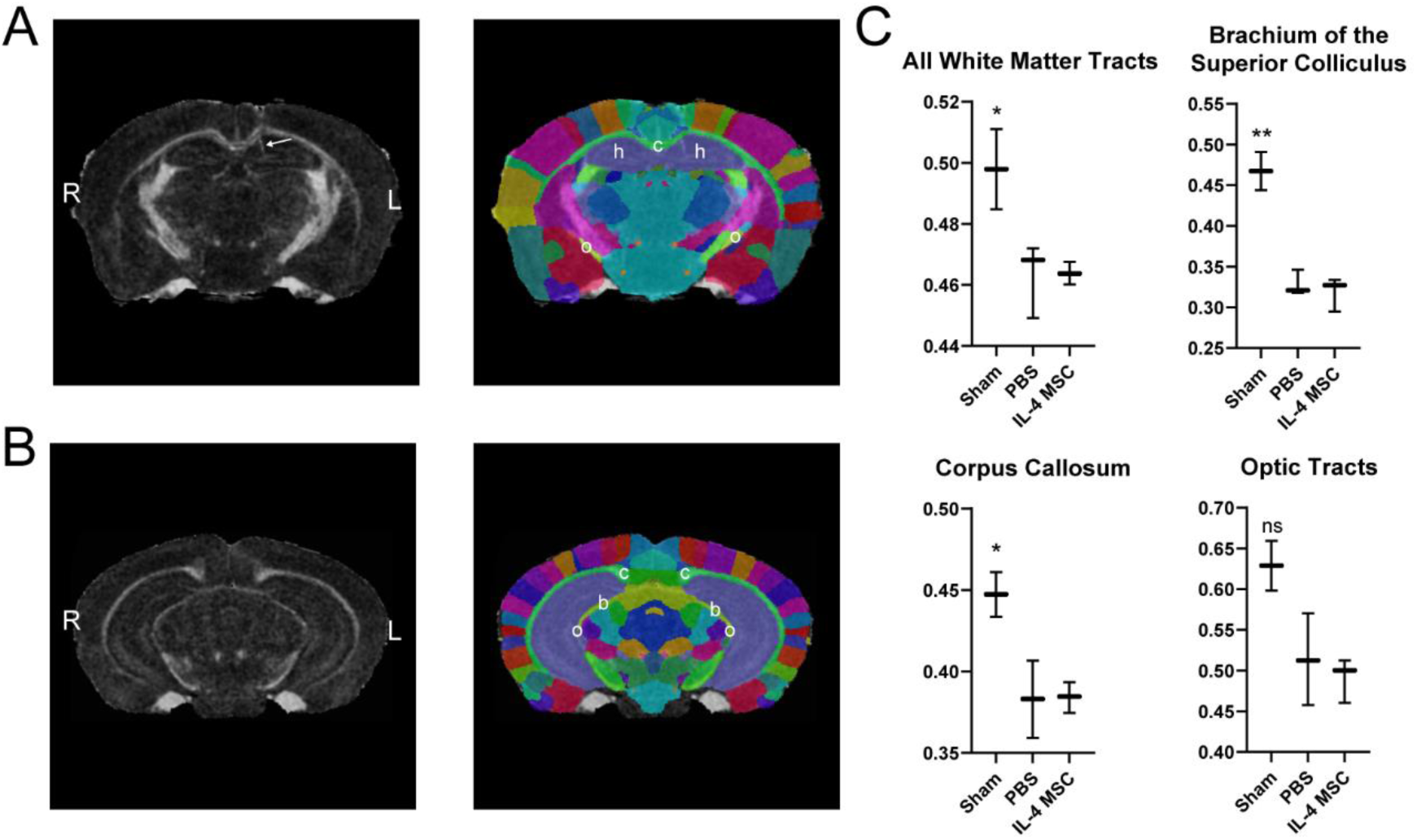
Diffusion-weighted magnetic resonance imaging to assess white matter integrity one month after injury. **(A), (B)** Left images are representative coronal brain slices of fractional anisotropy. Right images are corresponding map of cortical regions. Arrow indicates injection artefact. b = brachium of the superior colliculus; c = corpus callosum; h = hippocampus; o = optic tract. **(C)** Fractional anistropy values of each region per treatment group. Sham *n* = 2, PBS *n* = 3, IL-4 MSCs *n* = 3, N = 8; graphs present mean ± SD; one-way ANOVAs with post-hoc Dunnett’s multiple comparison against the PBS group; ns = not significant, \**p* < 0.05, \*\**p* < 0.01.

### i. Transcriptomics elucidates persistent inflammation and poor regeneration

To develop a mechanistic understanding of why elevating M2-like macrophage counts acutely did not improve most measures of healing, we investigated the transcriptome at the site of treatment. Raw transcriptomic data is accessible via the NIH Gene Expression Omnibus (GSE144193) and analyzed data are included as supplements to this paper. These results (Fig. 9) highlight selected findings relevant to our search for mechanisms that may have hindered healing. One sample was excluded due to anomalously different transcript counts (89-5, a PBS.3WK replicate). Overall, IL-4 MSCs had more significantly upregulated genes at 1-week post-injury than any other group. This was absent at 3 weeks (Fig. 9A).

**Figure 9:**
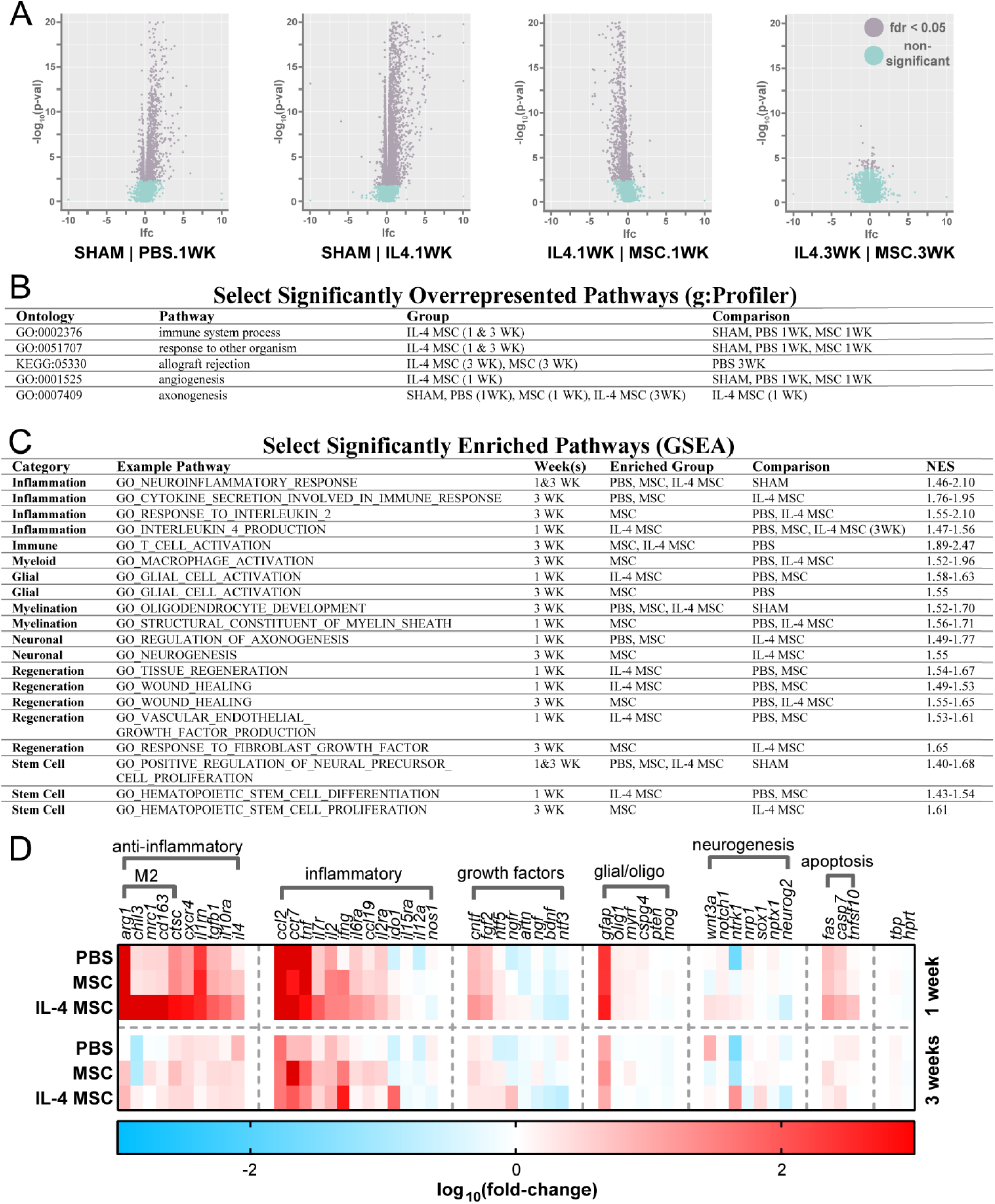
Transcriptomic analysis of mouse brains local to site of intervention at 1 and 3 weeks. **(A)** Volcano plots demonstrating that IL-4 MSCs induced the most alterations in gene expression. Each graph presents genes of the latter condition vs. the former condition. **(B)** Select significantly overrepresented pathways identified through g:Profiler. **(C)** Select significantly enriched pathways identified through Gene Set Enrichment Analysis and Molecular Signatures Database v7.0, Collection C5 **(D)** A heatmap presenting a selection of genes up- or down-regulated compared to Sham group.

As expected, numerous immune response and inflammation pathways were upregulated acutely in all groups compared to sham. However, although leukocyte numbers and inflammatory cytokines were reduced in prior data (Fig. 4 and 6), inflammation-associated pathways were still upregulated at 3 weeks. Specifically, cytokine, inflammatory interleukin, T cell, and macrophage pathways were chronically upregulated by MSCs compared to PBS, but this effect was absent in the IL-4 MSC group. Acutely, IL-4-associated pathways were enriched in the IL-4 MSC group compared to PBS and MSCs. However, these pathways were not enriched at 3 weeks, which suggests the IL-4 response does not persist. Thus, while these inflammatory pathways were over-represented in the MSC group compared to IL-4 MSCs, general inflammation pathways was chronically enriched in all groups compared to sham.

Alongside persistent inflammation, pathways of axonogenesis were over-represented and enriched in the sham group compared to all other groups at both acute and chronic time-points. However, axonogenesis pathways were also acutely overrepresented in MSC and PBS groups compared to IL-4 MSCs. One of these, “GO_NEGATIVE_REGULATION_OF_AXONOGENESIS”, also suggest down-regulation of axonogenesis in both groups relative to IL-4 MSCs. At the 3-week timepoint, the pathway, “GO_NEUROGENESIS”, was enriched in MSCs compared to IL-4 MSCs. Regarding myelination, few related pathways were enriched among injury and treatment groups, but at 3 weeks oligodendrocyte pathways were enriched compared to sham. Glial pathways were found to be enriched acutely for IL-4 MSCs, and in contrast, chronically for MSCs.

Acutely, IL-4 MSCs exhibited enrichment of regeneration, healing, stem cell, and angiogenesis pathways relative to MSCs and PBS. However, at the 3-week timepoint, MSCs tended to have higher normalized enrichment scores for these pathways compared to IL-4 MSCs. IL-4 MSCs also exhibited more enrichment of endothelial and platelet derived growth factor pathways at acute timepoints, while MSCs showed enrichment for fibroblast and insulin-like growth factor pathways, chronically.

All injury groups elicited enrichment for “response to other organism”, suggesting similar neural responses between injury and infection. PBS at 3 weeks also had over-representation for neurofibrillary tangles and tau protein pathways compared to sham, suggesting some overlap in TBI response to that seen in Alzheimer’s Disease. At 3 weeks, IL-4 MSCs and MSCs had over-represented pathways suggesting autoimmune activation, including graft-versus-host-disease and allograft rejection.

To begin to understand effects on microglia, we performed over-representation analysis of our gene data against a database that clustered microglia genes based on different stimulation conditions (S. Table 1). As expected, a cluster representing IL-4 stimulation of microglia (‘SA_Salmon’) was over-represented in the IL-4 MSC group at 1 week compared to both PBS and MSC. However, a few other clusters representing IFN and LPS stimulation were also over-represented at the 1-week timepoint. At the 3-week timepoint, both MSCs and IL-4 MSCs had over-represented clusters representing IFN and LPS stimulation compared to sham (BR_Blue, BR_Turquoise, PI_Turquoise). However, among the injury groups there were little differences. IL-4 MSCs had one cluster over-represented compared to PBS (BR_Turquoise) that suggested a response analogous to persistent IFNγ and LPS stimulation.

Overall, our analyses suggest that: 1) acute benefits of IL-4 MSCs do not persist; 2) inflammation persists to at least 3 weeks even with treatment; and 3) neural regeneration does not increase with treatment.

## C. DISCUSSION

One important reason a patient suffers disability after TBI is because the immunological response becomes maladaptive. To shift the immune response towards a regenerative phenotype, clinical trials are investigating therapeutic stem cell delivery. However, these stem cells do not secrete some potent regulators of inflammation. One absentee, IL-4, is a strong inducer of the anti-inflammatory, M2-like phenotype in macrophages and microglia. According to the literature, M2-like macrophages are associated with improved biological and functional outcomes in pre-clinical TBI models. Thus, in this study, we augmented MSCs to transiently express IL-4 and consequently observed a robust increase in M2-like macrophages and some anti-inflammatory genes after TBI. Curiously, this biological change did not improve functional or histological outcomes, though our transcriptomic analyses provide some potential answers as to why not.

We began this work by we transiently expressing IL-4 from MSCs and studying *in vivo* IL-4 levels and M2-like macrophage polarization. IL-4 has been well-documented as an inducer of this phenotype in macrophages and microglia *in vivo* in CNS and PNS injury models. In fact, transfection of macrophages with IL-4 mRNA also promotes an M2 phenotype *in vitro* and *in vivo* (84). Our results corroborated this as we saw a dramatic rise in the polarization and quantity of M2-like macrophages at both 2 and 7 days after our intervention (about 1 and 2 weeks after injury). Simultaneously, we witnessed upregulation of numerous anti-inflammatory and M2-like genes. However, while this polarization may be desirable, IL-4 induces macrophage proliferation as well. Thus, it might be important to understand how much IL-4 is required *in vivo* and how many M2-like macrophages are optimal for neuroprotection. Our data begin to answer some of these questions. We saw that a bolus of recombinant IL-4 is less effective at polarizing macrophages than sustained expression over 24-48 hours from IL-4 MSCs (Fig. 4). Separately, we observed that a small proportion of IL-4 from IL-4 MSCs reaches the contralateral hemisphere (Fig. 6A). Surprisingly, this induced equivalent polarization to the ipsilateral hemisphere (S. Fig. 3F). Critically, however, the total number of macrophages and specifically M2-like macrophages was significantly lower in the contralateral hemisphere (S. Fig. 3 E and G). Finally, 1 week after intervention, IL-4 MSCs still had a greater number of M2-like macrophages compared to other groups, though the polarization percentage had dropped. Most of the macrophages did not express CD206 or CD86, suggesting they may possess a phenotype we did not explore. Our transcriptomic data also demonstrated a signature for persistent immune and inflammatory response in all injured groups compared to sham. Furthermore, IL-4 response pathways did not persist with IL-4 MSC treatment. Thus, hypothetically, an ideal scenario would recruit much fewer macrophages, with the majority biased to be M2-like. These data suggest that a sustained yet smaller dose of endogenous IL-4 could foster that.

Second, we tested our IL-4 MSCs in a TBI model of CHI but did not see an acute improvement in motor coordination or memory. Numerous CHI models exist, and these have been extensively reviewed (85). We believe that since CHI induces a diffuse injury, it is more clinically relevant than focal TBI models. Studying behavioral outcomes, however, can be tricky. The central understanding is, if function is improved, the therapy had long-reaching effects; it not only influenced biology directly, but ultimately promoted neuroprotection, regeneration, or plasticity. However, TBI models have high intrinsic biological and technical variability. Additionally, behavioral experiments can be plagued with invisible biases. Thus, for all TBI studies assessing behavior, it becomes critical to: 1) demonstrate that injury induced equivalent dysfunction in all groups; and 2) use statistics that correct for repeated measures. Our injury induced a significant behavioral deficit (Fig. 5 B and C), but IL-4 MSCs did not rescue Rotarod performance or memory (Fig. 5D-F). Unfortunately, this corroborates many other attempts with genetically modified MSCs (49,63,65,67). Studies that saw an improvement typically saw a change in only one out of multiple assessments, or only looked at a single time point (63,64,66). In one study delivering recombinant IL-4 after stroke in IL-4 knockout mice, repeated measures were not accounted for and wild-type mice were not tested (52). A variety of other approaches to TBI that promote an M2-like response have also observed improvement in behavioral outcomes (21–30) (31–36). However, in many of these studies, behavior is not (or could not be) assessed prior to treatment to ensure the absence of accidental bias. Assessments of righting time and NSS may be of use, but our data suggest that these metrics are not predictive of behavioral performance (S. Fig. 4). In this study we also only looked at acute and sub-acute timepoints. It is conceivable that the M2-like macrophages might have limited the spread of damage. Benefits of this might only be seen over months as neural rewiring slowly takes place. Interestingly, our transcriptomic analysis of IL-4 MSCs demonstrated that pathways relating to regeneration and healing were enriched at the 1 week time point. However, pathways more relevant to neural function (e.g. axonogenesis, neurogenesis) were overrepresented or enriched in the other groups instead. Thus, our data suggest that either our method of IL-4 MSC intervention alone cannot alter function, or that our behavioral assessments may not have been adequately sensitive to reveal the effects of IL-4 MSCs on TBI.

Third, while injury alone induced histological changes, we observed damage caused by intrahippocampal injections as well. Histology demonstrated significant increase in GFAP across all injured groups. GFAP is a marker for reactive astrocytosis, an endogenous response to limit deleterious effects of TBI. However, chronically, astrocytosis also prevents neural regeneration. Interestingly, transcriptomic analysis suggested that gliosis pathways were only enriched acutely in the IL-4 MSC group compared to PBS and MSC groups, a potentially desirable scenario. At the 3-week time point, it was MSCs that had gliosis pathways enriched compared to IL-4 MSCs. However, both the amount of gliosis observed histologically and the transcriptomic enrichment of gliosis pathways compared to sham suggest that IL-4 MSCs do not reduce gliosis in a biologically relevant manner within 3 weeks. IL-4 MSCs did, however, increase the number of macrophages and microglia (indicated by CD68 staining and corroborated by flow cytometry). However, the needle used to inject the treatments, and possibly the bolus of cells, negatively impacted the architecture of the ipsilateral hippocampus. This likely significantly reduced memory performance in the MWM task (Fig. 5E). We chose intrahippocampal delivery over intraventricular due to the improved polarization (S. Fig. 3F). Additionally, one TBI clinical trial is exploring stereotactic intracerebral delivery of modified MSCs (NCT02416492). However, an intraventricular delivery would have saved the hippocampus and likely left a smaller cortical injury as the cerebral spinal fluid and ventricles can better accommodate a transiently increased volume. In our studies, we did not assess intravascular delivery as <0.1% of MSCs cross the BBB (86) even though it may be permeable soon after TBI (87). However, this may still be a viable approach. In this case, instead of targeting IL-4 MSCs toward the brain, the modified MSC secretome could influence peripheral monocytes prior to invading the TBI environment. In fact, many of the TBI interventions that improved behavioral outcomes were through intravascular routes. Interestingly, IL-10 improved outcomes when administered intravenously but not when delivered intracranially (88). This difference in effect has also been observed with cord blood cells for stroke (89). Thus, our data and the literature suggest that systemic delivery may be the better avenue for TBI even though the BBB is generally considered an obstacle.

Finally, our studies also suggest that CD206^+^, CD86^-^ macrophages alone may not be able to influence all the other maladaptive processes after TBI. There are conflicting reports of whether adoptive transfer of M2 macrophages can promote healing after CNS injury (27, 90). The M2 phenotype can be further divided into a, b, and c subtypes (91). M2a phenotypes are promoted by IL-4 and IL-13 and promote tissue repair and stimulate growth. This is the phenotype we likely enriched and explains the large increase in CD206^+^ macrophages as well. M2b macrophages can be proliferative and secrete a mixture of inflammatory and anti-inflammatory cytokines. M2c phenotypes are induced by IL-10, can promote fibrosis, but are also known to be deactivated. In spinal cord injury, the phenotypes follow each other sequentially with an M2a peak 1-3 days after injury and M2b or M2c peak 4-6 days after. We do not know whether this holds true in TBI; only that M2-like macrophages may peak around day 5 (37, 38) or even day 7 (92) but are quickly overpowered by M1 phenotypes (9). We also do not know whether the endogenous pattern of sequential phenotypes is ideal for regeneration in neural organs. Getting the right subtype of macrophage, at the right time, may be critical to reducing inflammation and promoting homeostasis. In these subtypes, there might be an explanation for why delivering IL-4 or IL-10 alone has been ineffective. Perhaps, by appropriately timing each cytokine in a concerted approach, M2-like macrophages could proliferate early to overpower maladaptive processes, and then be deactivated as desired. Additionally, we did not scrutinize microglia in our flow cytometry studies as they do not reliably express CD206 (93). Distinguishing macrophages from microglia is not trivial but IL-4 is known to have favorable effects on both (58,94,95). However, as M1 and M2 phenotypes concurrently exist on microglia, they may need to be assessed for a favorable phenotype after an IL-4 MSC intervention (13). To obtain some clues to how IL-4 MSCs were affecting microglia, we compared our transcriptomic signatures against a published microglia transcriptomic database. Even with the caveat that our transcripts were obtained from bulk tissue and not isolated microglia, we saw overrepresentation of microglial ‘clusters’ that suggest a signature of microglia responding to IL-4 only in the IL-4 MSC group. However, alongside this, all injury groups had clusters suggesting similarities to IFN and LPS stimulation of microglia as well. Therefore, like the macrophages, microglia appear to be receiving numerous signals and IL-4 alone is not enough to adequately modulate the biology.

These studies demonstrate a method to augment MSCs with synthetic IL-4 mRNA, a modified closed head injury model of TBI with a behavioral deficit, and also explore the effect of acute IL-4 and M2-like macrophage enrichment after TBI. By inducing MSCs to produce IL-4, we also circumvent the challenges of recombinant protein delivery. Moving forward, it will be useful to explore the effects of dosing, dual-cytokine enrichment, and intervening systemically. Lower doses of either IL-4 mRNA or IL-4 MSCs may promote better healing; although a greater dose may be required for intravenous administration. A few dual cytokine strategies could include: timed deliveries of IL-4 MSCs and IL-10 MSCs; a single delivery of MSCs transfected with IL-4 mRNA (rapid expression) and IL-10 DNA (delayed expression); or intravenous IL-10 bolus a few days after IL-4 MSCs. Additionally, alternate stem cells could be transfected with synthetic IL-4 mRNA, such as MAPCs and umbilical cord cells which may be more effective in TBI than MSCs (40, 44). Alternatively, MSC exosomes possess benefits over MSCs themselves and could be augmented through transfection (96, 97). These approaches may also have merit outside TBI, as IL-4 MSCs may facilitate healing in any wounds or environments where proliferation is desired, e.g. in cavities left from disease, or to facilitate tissue integration with implanted biomaterials (17).

Beyond alternative therapeutic approaches, the question remains of how to best modulate inflammation to improve outcomes in TBI. The answer requires a deeper understanding of the roles of macrophage and microglia sub-phenotypes and their interaction with native cells in a TBI environment. Importantly, we need to develop a better understanding of what cytokines and cells are required and when should they be enriched after TBI. As our transcriptomic data suggested that infection and injury have similar pathway enrichment, models of alternate inflammatory stimuli, in alternate orgniams, or even an evolution-centered inquiry may provide insight into the maladaptive immunological response seen after TBI (98).

In conclusion, MSCs that transiently express IL-4 can robustly polarize macrophages toward an M2-like phenotype for at least one week after TBI. While this induces cytokine and gene-level changes, it alone is not enough to significantly impact histological or functional outcomes. Transcriptomic studies reveal persistent inflammatory pathways and an absence of neuroregeneration. However, the question of whether this is due to insufficient or excessive levels of IL-4 or M2-like macrophages, a lack of sensitivity in histological and behavioral measures, or the ineffectiveness of a single cytokine or cell phenotype in a complex environment, remains.

## D. MATERIALS AND METHODS

### a. *In vitro* transcribed mRNA (IL-4, Luciferase, GFP)

All synthetic mRNA were *in vitro* transcribed in the Santangelo Lab (Georgia Institute of Technology, US) similarly to past research (84).

## b. Cells (MSCs, Macrophages)

Mesenchymal stem/stromal cells (MSCs) were purchased at passage 6 from Cyagen Biosciences (US) (Lot#: 170221|31). These MSCs are primary cells harvested by the vendor from C57BL/6N mice. All *in vivo* experiments used the same strain of mice. According to the vendor, the MSCs express CD29, CD44, and Sca-1 and do not express CD31 and CD45. We verified this expression in two late passages (S. Fig. 1). The MSCs were grown in Mouse Mesenchymal Growth Media (Cyagen Biosciences, US) with FBS (Lot# T161102G002) and frozen at Passage 7 and stored in the vapor phase of liquid nitrogen. For every *in vivo* experiment, the cells were thawed, and grown in media for 3-4 days before experiments. To passage or plate cells, the cells were washed twice with PBS (Corning, US). Then 0.25% Trypsin (Corning, US) was added, with enough volume to cover the surface area, and gently rocked for 90s. The Trypsin was suctioned off, and the cells were left in a 37°C incubator for 1 minute. Finally, the cells were harvested by adding fresh, pre-warmed, and pH-equilibrated media to the cells.

For macrophage polarization experiments, J774A.1 macrophages were obtained from the Cell Culture Facility at Duke University (thawed from an unknown passage number). These were grown in DMEM-High Glucose (Gibco, US) supplemented with 10% FBS. Cells were passaged by manual dissociation with a cell scraper.

### c. Transfection

Viromer Red (Lipocalyx, US), jetPEI (Polyplus Transfection, US), and Lipofectamine Messenger Max (ThermoFisher, US) were used to transfect MSCs with GFP mRNA following manufacturer protocol. All subsequent MSC transfections of synthetic IL-4 or Luciferase mRNA used Viromer Red, following its protocol. Prior to every transfection, a cell-culture cabinet, pipettes, and other materials were decontaminated with 70% ethanol and RNAse Away (ThermoFisher, US) and left to dry.

### d. *In vitro* flow cytometry

For MSC cell-surface characterization, the following antibodies were used: APC-Vio770 CD29 (Miltenyi Biotec; clone: HMβ1-1), VioBright FITC CD44 (Miltenyi Biotec, US; clone: REA665), PerCP Sca-1 (Biolegend, US; clone: D7), PE CD31 (Miltenyi Biotec, US; clone: 390), PE CD45 (Biolegend, US; clone: 30-F11), and PE-Vio770 CD105 (Miltenyi Biotec, US; clone: MJ7/19). For analysis of *in vitro* macrophages we first blocked Fc receptors with: CD16/32 FcγRIII/FcγRII (BD Bioscience, US; clone: 2.4G2) and CD16.2 FcγRIV (Biolegend, US; clone: 9E9) and then stained with: PE CD45 (Biolegend, US; clone: 30-F11), Alexa-Fluor 647 CD206 (BioLegend, US; clone: C068C2) and PE CD163 (ThermoFisher, US; clone: TNKUPJ). A Novocyte 2060 (ACEA Biosciences, US) was used to measure mean fluorescence intensity of GFP from transfected MSCs, MSC cell-surface markers, and macrophage markers. Data was analyzed in FlowJo software (BD Biosciences, US).

### e. *In vitro* assays: viability, ELISA, imaging

Cell viability was assessed with either a Live/Dead Viability/Cytotoxicity kit (ThermoFisher, US) and DMi8 LiveCell Microscope (Leica Biosystems, US) or 0.4% Trypan Blue (ThermoFisher, US) exclusion with automated cell counting using a calibrated Countess II (ThermoFisher, US). For protein quantification of conditioned media, an IL-4 colorimetric ELISA was performed (R&D DuoSet; R&D Systems, US) with appropriate ancillary kits. Optical density was obtained via a 96-well plate reader (SpectraMax i3x, Molecular Devices, US). In some experiments, prior to media collection, cells were imaged under brightfield in the DMi8 LiveCell Microscope and the images were analyzed with a custom ImageJ script to quantify cell coverage of the well in each well as an estimator of cell confluency/count.

### f. Closed head injury

All *in vivo* experiments were conducted according to protocols approved by the Duke Institute for Animal Care and Use Committee (IACUC). Male C57BL/6N mice were purchased from Charles River Laboratories, US and left to acclimate for 1-2 weeks in Duke University animal housing. Procedures began when the mice were between 8-9 weeks of age.

Closed head injury (CHI) experiments were modified from *Webster et. al.,* to develop a model that is easily reproducible but also induces behavioral deficits in motor coordination (77). Mice were induced under 5% isoflurane vapor anesthesia and maintained between 1-3% anesthesia. Hair was trimmed from the scalp and ointment was applied on the eyes (Paralube; Dechra, US). The scalp was sterilized with three 70% ethanol and chlorhexidine swabs each. A mid-line cut into the scalp was made with micro-scissors, the cranium was exposed, and the bone surface cleaned with a cotton-tip swab. The mouse was then transferred onto a custom heated bed in a rat stereotactic apparatus (51600; Stoelting, US) with mouse gas adaptor (10030-386; VWR, US), and mouse non-rupture ear bars (922; Kopf Instruments, US) connected to a separate anesthesia system maintaining 1-3% anesthesia. A balloon connected through tubing to a 20-mL syringe was also placed under the head of the mouse to prevent skull fracture (77). A 5-mm impact probe connected to an electromagnetic impact system (Impact One; Leica Biosystems, US) was extended and positioned −1.5 mm AP from bregma and zeroed to the surface. At this point anesthesia was reduced to 0.5%, oxygen flow increased to 2 L/min, and the impactor retracted and then lowered 1.5 mm DV. When breathing became faster, a toe-pinch was assessed to ensure no pain sensation, and the impact was hit at 6 m/s. If the toe-pinch induced a reflex, injury was not performed, anesthesia was immediately increased, and the steps were repeated. Controlling anesthesia levels carefully is critical to survival; if the anesthesia is too deep prior to impact, the mouse may not return to spontaneous breathing after the impact. Immediately after impact, the oxygen flow was increased to 4 L/min and the mouse was placed on its right side for chest compressions carried out with an index finger and thumb. When the mouse began to breathe spontaneously, the scalp incision was stapled (Reflex 7-mm wound clips, Roboz Surgical, US), 3 drops of 0.5% bupivicaine were added to the site, and the mouse placed supine in an empty cage. Time for the mouse to right itself into a prone position was recorded. Righting was considered when all four paws were on the cage floor. If there was a fracture, brain hernia, or excessive bleeding after the impact, the mouse was immediately put back under 5% anesthesia and then euthanized. If righting time was >10 minutes, it was considered an injury (79). Neurologic Severity Score (NSS) was assessed in two experiments as per its protocol (80). With two scientists working on mouse preparation and injury separately, injuries can be 5-10 minutes apart. Sham procedures consisted of all steps except for the impact.

### g. Preparation and delivery of MSCs

Based on transfection and expression data (Fig. 3A), an mRNA incubation time of 10 hours was chosen. On the day prior to injections, cultured MSCs were plated in the afternoon in 6-well plates at 300,000 cells per well. After at least 6 hours, MSCs were transfected with 2 µg mRNA complexed with Viromer Red according to manufacturer protocol. Ten hours after transfection, the MSCs were harvested with two 1-mL collections of media per well. All similar wells were grouped in 15 mL tubes and spun down at 300xg for 3 minutes. The media was suctioned, and the cells were resuspended in 1 mL of PBS. The cells were counted via a cell counter (Countess II; ThermoFisher, US) that was previously calibrated to manual cell counting. The cells were then spun again and resuspended to make a 30 million cells/mL mixture in PBS. Aliquots of 10 µL were made and kept on ice until injection.

To deliver MSCs, intrahippocampal injections were conducted two or five days after injury. As previously, the mice were induced with 5% isoflurane and maintained between 1-3% anesthesia. The eyes were protected with ointment, the surgical site was cleaned with ethanol and chlorhexidine, staples were removed with a staple remover, the old incision was opened with micro-scissors, and the skull surface was cleaned with a cotton-tip swab. The mouse was moved to the rat stereotactic apparatus with a different mouse gas adaptor (923-B; Kopf Instruments, US). A craniotomy was performed with a 0.6-mm drill-bit (Roboz Surgical, US) attached to a handheld drill (Stoelting, US) at −1.5 mm AP, and −1 mm ML (left) drilling 0.4-0.6 mm deep. Cells were then mixed and picked up by a 5 µL syringe (75RN; Hamilton, US) with a 26G needle (1-inch, point style 4, 30°; Hamilton, US). This needle was chosen as it had the closest inner-diameter to the syringe. The syringe was attached to the stereotactic apparatus and inserted to a depth of 2 mm DV from the outer surface and then retracted 0.5 mm for a final injection location of 1.5 mm DV (left hippocampus). For ventricular injections the coordinates were −0.5 AP, −1 ML, 2.25 DV. Cells were injected at a rate of 0.5 µL/min in a volume of 5 µL.

### h. Behavioral experiments (Rotarod, Morris Water Maze)

To assess motor coordination and function, mice were tested on an accelerating Rotarod apparatus (Med Associates Inc, US). On the day prior to CHI, the mice first underwent a training trial on the rotarod where they were put back on as they fell. The rotarod accelerated from 4 to 40 rotations per minute. After that, three recording trials were run with at least 15 minutes between each trial. Mice were removed if they stayed on for 400s or if they held onto the rod without running for two spins. The time on the rotarod was recorded.

To assess spatial memory function, a Morris Water Maze was set up with a large round pool, four images on the walls, a glass platform hidden under the surface of the water, and a camera and computer set up with tracking software (ANY-maze; Stoelting, US). Mice were placed into the water starting at the edge of the pool and time to find the hidden platform was recorded. Four trials were conducted each day, each starting at one pole of the pool (North, South, East, or West).

### i. Perfusion/euthanasia

All mice were sacrificed under 5% isoflurane anesthesia and cardiac perfusion. Perfusate consisted of Hank’s Balanced Salt Solution (Corning, US) with Heparin (10,000 units/L; ThermoFisher, US) kept on ice. After testing for a reaction to toe-pinch, the sternum was lifted, and the abdomen and ribs cut with scissors. The diaphragm was dissected away, and the heart was exposed and freed from connective tissue. The left ventricle of the heart was pierced with a 26G 3/8” needle (BD Biosciences, US) and the mouse perfused at a rate of 8 mL/min for at least 2.5 minutes with a motorized pump (Cole-Parmer, US). For immunohistochemistry, this was followed by cold 4% formaldehyde in HBSS. The mouse was then decapitated, and the brain harvested. For protein and gene expression studies, the brain was further dissected to isolate the injection region (3-mm width per hemisphere), stored in 5 mL conical tubes, frozen in liquid nitrogen, and stored in a −80°C freezer until processed.

### j. *Ex vivo* flow cytometry

To study macrophage polarization, leukocytes were isolated from freshly harvested brain tissue following previously published protocol (99). The protocol was slightly modified and is included in the supplementary information. As previously, Fc receptors were blocked with antibodies against CD16/32 FcγRIII/FcγRII (BD Bioscience, US; clone: 2.4G2) and CD16.2 FcγRIV (Biolegend, US; clone: 9E9) and then stained with PE-Cy7 CD45 (BioLegend, US; clone: I3/2.3), PE F4/80 (BioLegend, US; clone: BM8), Alexa Fluor 488 CD86 (BioLegend, US; clone: GL-1), and Alexa Fluor 647 CD206 (BioLegend, US; clone: C068C2). Macrophages were defined as CD45^high^, F4/80^+^. Cells were run in a Novocyte 2060 (ACEA Biosciences, US) at a rate of 14 µL/min.

### k. *Ex vivo* protein quantification

Frozen brain tissue samples were homogenized in their tubes with a TissueRuptor II (6 speed, 30s; Qiagen, US) and disposable probes (Qiagen, US) on ice with Halt Protease Inhibitor (ThermoFisher, US) in N-PER reagent (ThermoFisher, US). Homogenates were spun down thrice. The first spin in the 5 mL tubes was at 1000xg for 10 minutes at 4°C. The supernatant was transferred into 2 mL tubes. The second and third spins were at 12,000xg for 15 min and 15,000xg for 20 min respectively, both at 4°C, with a transfer to new tubes after each spin. The protein samples were then processed with an 8-plex bead-based assay (LEGENDplex Mouse Th1/Th2 Panel; BioLegend, US) according to manufacturer protocol. Samples were run on a Novocyte 2060 (ACEA Biosciences, US). Quantification was normalized against a total protein bicinchoninic acid (BCA) assay (ThermoFisher, US) of each protein sample.

### l. Gene expression and RNA-Sequencing

The workspace was first cleaned and sprayed with RNAse Away (ThermoFisher, US). Frozen brain tissue samples were processed with an RNeasy Lipid Tissue Kit (Qiagen, US) to extract RNA. The tissue samples were homogenized with a TissueRuptor II (6 speed, 30s) and disposable probes. RNA integrity was assessed on an Agilent Bioanalyzer (Agilent, US) with the assistance of the Duke Microbiome Shared Resource. Complementary DNA (cDNA) was synthesized with SuperScript IV VILO (ThermoFisher, US), separated into aliquots and stored at −80°C. Genes were studied with 26 Taqman Probes (ThermoFisher, US) in 384-well plates. The samples were run on a 7900HT Real-Time PCR system (Applied Biosystems, US). Data was analyzed using Applied Biosystems Analysis Software available on the ThermoFisher Cloud Connect Utility. Global normalization was used, and Sham groups were kept as the reference for each gene.

For RNA-Sequencing, total RNA was sequenced through Illumina NovaSeq 6000 (50 bp, paired-ends). The raw data was quality-checked, adaptors were trimmed with cutadapt (v2.4), and then pseudo-aligned using Kallisto (v0.46.0) against protein-coding GENCODE transcripts (vM23)(100), with differential expression assessment via Sleuth (v0.30.0) in R (v3.6.1) (101, 102). Data were consolidated and organized using Python (v3.6.5, w/ pandas 0.24.2, numpy 1.16.4) and then run through an over-representation analysis (g:Profiler, rev 2fcb244) and Gene Set Enrichment Analysis (GSEA, v4.0.3) (103, 104).

### m. Immunohistochemistry

To study histological changes after intervention, 4% formaldehyde-fixed brains were embedded in optimal cutting temperature compound (Tissue-Tek; VWR, US) and frozen in a bath of 4-methylbutane chilled in liquid nitrogen. The brains were sectioned in 10-µm slices and collected on Superfrost Plus slides (ThermoFisher, US) with a hydrophobic barrier drawn (ImmEdge Hydrophobic Barrier PAP Pen; Vector Laboratories, US). Slides were kept at −20°C until ready to be stained. Primary antibodies included: Chicken anti-mouse GFAP (abcam, US), rabbit anti-mouse Iba1 (abcam, US), and rat anti-mouse CD68 (abcam, US). Secondaries included: DyLight 488 goat anti-chicken IgY (abcam, US), Alexa Fluor 594 goat anti-rat IgG, and Alexa Fluor 680 goat anti-rabbit IgG (abcam, US). Cell nuclei were counterstained with DAPI (Sigma-Aldrich, US). Coverslips were applied after drops of Fluoromount-G (SouthernBiotech, US) were added. One group of slides was stained with a Fluoro-jade C kit (Biosensis, US). Coverslips were applied after drops of DPX Mounting Media (Sigma-Aldrich, US) were added. All staining procedures were conducted at the same time for each group. Imaging was done in a DMi8 microscope (Leica Biosystems, US). All images were acquired in the same time period per stain. Images were analyzed with a custom semi-automated script on ImageJ with regions of interest drawn for hippocampi, cortices, and corpus callosum.

### n. MRI

Diffusion-weighted magnetic resonance imaging was conducted with the Duke Center for In Vivo Microscopy (CIVM). MR images were acquired on a 7.0T horizonal bore magnet with Resonance Research gradients providing ∼650 mT/m maximum gradient. The system is controlled by an Agilent Direct Drive console. Specimens were mounted in a 12 mm diameter single sheet solenoid radiofrequency coil. Three-dimensional (3D) diffusion weighted images were acquired with a Stejskal Tanner rf refocused spin echo sequences with TR/TE of 100/21.15 ms and b values of 4000 s/mm^2^. Compressed sensing was used with an acceleration of 8X to reduce the acquisition time (105). The result is a 4D image array with isotropic spatial resolution of 45 um (voxel volume of 91 pl).

The 4D array (256×256×420×7) was passed to a post processing pipeline that registered each of the diffusion weighted 3D volumes to the baseline to correct for eddy current distortion. The registered 4D array was passed to DSI Studio which generated scalar images (AD, RD, FA, ClrFA) using the DTI model. A 3D label set was registered onto each 4D volume in that volume’s native reference frame i.e. the reference frame in which the data were acquired. Waxholm Space (WHS) (106) an isotropic label set for the mouse brain was expanded by Calabrese et al was extended with an additional 18 regions of interest yielding 166 regions of interest on each half of the brain (107).

### o. Statistics and graphing

All statistics were performed in Prism V8.2 (GraphPad, US). Where appropriate, Student’s t-test, one-way ANOVAs, repeated measures ANOVAs, and two-way ANOVAs were carried out. All data are presented as means ± standard deviation. Graphs were made in Prism V8.2 and converted into figures in Photoshop and Illustrator (Adobe, US).

## Acknowledgements

We are grateful for the invaluable help of: Jared Beyersdorf and Jonathan Kirschman of the Santangelo Lab at the Georgia Institute of Technology for synthesizing IL-4 mRNA *in vitro*; G. Allan Johnson, Gary Cofer, James J. Cook, and the Center for In Vivo Microscopy at Duke University for magnetic resonance imaging and mapping; and Nicholas Devos, David Corcoran, Heather Hemric, and the Duke Center for Genomic and Computational Biology for RNA and transcriptomic analysis.

